# An ultra-fast, proteome-wide response to the plant hormone auxin

**DOI:** 10.1101/2022.11.25.517949

**Authors:** Mark Roosjen, Andre Kuhn, Sumanth K. Mutte, Sjef Boeren, Pavel Krupar, Jasper Koehorst, Matyáš Fendrych, Jiří Friml, Dolf Weijers

## Abstract

The plant signaling molecule auxin controls growth and development through a simple nuclear pathway that regulates gene expresion. There are however several cellular and physiological responses to auxin that occur within seconds, far too rapid to be mediated by transcriptional changes, for which no molecular mechanism has yet been identified. Using a phosphoproteomic strategy in *Arabidopsis thaliana* roots, we identify an ultra-rapid auxin response system that targets over 1700 proteins, many within 30 seconds. Auxin response is chemically specific, requires known auxin-binding proteins, and targets various pathways. Through exploring its temporal dynamics, we infer auxin-triggered kinase-substrate networks and identify apoplastic pH changes as a target of signaling and as part of a relay mechanism. By generating a variety of phosphoproteomic datasets, integrated with structural information in a web-app, and by demonstrating analysis and inference strategies, we offer a resource to explore rapid and dynamic signaling in plants.

## INTRODUCTION

Since its discovery nearly a century ago^1,2^, the plant signaling molecule auxin has been the focus of intensive investigation^3^. The naturally occurring indole 3-acetic acid (IAA) is widespread in nature and found in essentially all plants studied to date^4^. Its biological activity as a signaling molecule, acting at nanomolar to micromolar concentrations, is extremely profound. When applied to plants or plant cells, auxin can trigger a wide range of physiological, cellular and molecular changes that likely underly the long-term effects on plant growth and development. Classical examples of auxin activity include mediating tropic growth (e.g. phototropism or gravitropism) ^1,2,5,6^, as well as the promotion of vascular cell differentiation^7–9^, and the induction of adventitious or lateral roots^10,11^. Molecular and cellular responses to auxin include rapid changes in membrane chemical potential, associated changes in ion transport^12–15^, changes in the velocity of cytoplasmic streaming^16,17^, changes in cellular growth rate and differentiation^11,18–20^, and importantly, reprogramming of transcription^21^.

To identify the mechanisms underlying the responses to auxin that control plant growth, genetic screens have led to the isolation of a range of Arabidopsis mutants that are insensitive to auxin’s growth-inhibiting action^22–24^. Cloning of the affected genes, followed by detailed molecular, biochemical and structural analysis have revealed a simple, yet powerful mechanism for auxin response^24^. In what is referred to as the Nuclear Auxin Pathway (NAP), auxin enables interaction of Aux/IAA transcriptional inhibitors with a ubiquitin ligase complex (SCF^TIR1/AFB^) to promote their degradation. This liberates DNA-binding ARF transcription factors from Aux/IAA inhibition and allows transcriptional control of their many target genes (reviewed in ^21^). This NAP can account for many of the observed auxin activities across land plants and in many developmental contexts, including responses to environmental signals^3^. However, the time required to bring about changes in cellular physiology through transcriptional control is substantial, and limited by the velocity of transcription, translation, protein folding and targeting. Hence, even if the first IAA-induced transcripts are visible between 5-10 minutes after IAA treatment^25^, it is extremely unlikely that the NAP can account for the changes in e.g. ion fluxes, cellular growth and subcellular traffic that have been observed to occur within seconds to minutes^12,14,16,17,26,27^. It is therefore inevitable that there exists a second auxin response system that is kinetically suited to mediate such rapid responses. Indeed, several “non-canonical” auxin responses have been reported^14,26,28^ that do not rely on a (complete) NAP, but that may involve one of its components. For example, fast inhibition of Arabidopsis root growth and changes in membrane polarity involve the AFB1 receptor of the NAP^14^. Other rapid responses including the activation of H^+^-ATPases at the plasma membrane for growth regulation^27,29^ and cytoplasmic streaming, possibly related functions in endocytic trafficking ^16,30^ require a cell surface auxin signaling mediated by the ABP1 auxin receptor and its associated TMK receptor-like kinase^3^. Thus, the existence of fast responses to auxin is evident, but it is unclear what mechanisms and response system(s) might underlie such fast responses.

Here, we explore the role that reversible protein phosphorylation may play in fast responses to auxin. Protein phosphorylation is a widespread mechanism for enzymatically modifying the structure and function of pre-existing proteins^31^, thus eliminating the need for *de novo* protein synthesis. Given that phosphorylation depends only on the (allosteric) activation of a protein kinase, the reaction is intrinsically rapid. Several well-known examples of phosphorylation-based signaling exist across the kingdoms of life^32–35^. Among these, some are particularly rapid, with Insulin and EGF ligands triggering initial phosphorylation changes by receptor kinases within seconds^34,35^, followed by relays and amplification steps with additional protein kinases^35^. Phosphorylation-based signaling is also widespread in plants, and mediates responses to peptide ligands in development and immunity^33^, as well as to Brassinosteroids^36–38^. Whether phosphorylation-based signaling is a meaningful component in auxin response is an open question. There is ample evidence for phosphorylation-based events in the control of activity and localization of PIN auxin transporters^39^. Whether however, these events are part of a rapid signaling system is an open question. The TMK1 receptor-like kinase mediates IAA-triggered phosphorylation changes, even within two minutes^16,27^, but the scope or dynamics of this response is unclear, given that the number of targets reported is limited.

Here, we used a phosphoproteomics strategy to address if a fast response system to IAA built on protein phosphorylation, exists. Prior phosphoproteomic analysis of auxin response has been reported in both Arabidopsis^27,40,41^ and rice^42^. In rice, a 3-hour treatment was reported ^42^, while in Arabidopsis, treatments used the synthetic auxin analogue 1-NAA for either 2 hours ^40^ or 11 days^41^, leaving the question of whether naturally occurring auxin can trigger phosphorylation changes at the same scale. We recently reported that IAA can trigger changes in protein phosphorylation in Arabidopsis roots within 2 minutes and that this response requires ABP1 and TMK1^27^. However, it is entirely unclear what the specificity, kinetics, scope and targets of this response are.

Here, we show that auxin triggers changes in the phosphorylation of hundreds of proteins in Arabidopsis roots well within 30 seconds. An extended time series showed that over two thousand proteins are differentially phosphorylated within 10 minutes, with a range of temporal dynamics. Analysis of mutants suggests these responses to involve both intracellular and extracellular auxin perception. We leveraged the rich datasets to identify pathways and functions that are triggered by rapid auxin signaling, predict signaling hub kinases and demonstrate a very rapid effect on apoplastic pH. Lastly, we compiled the datasets in a searchable web tool, integrated with protein structural information. We thus identified a novel layer in auxin response with profound impact on many cellular pathways and functions, and created a resource that will form the starting point for renewed mechanistic investigation in the underexplored area of fast auxin response.

## RESULTS

### Identification of a specific, rapid, phosphorylation-based auxin response

To explore the ability of Arabidopsis roots to respond to auxin by changing protein phosphorylation levels, we initially treated roots with a concentration (100 nM) of the naturally occurring auxin Indole 3-Acetic Acid (IAA) that is sub-maximal with regards to physiological responses. The earliest recorded transcriptional responses to auxin treatment occur after 5-10 minutes^25,43^. Also given theoretical considerations related to gene expression and protein synthesis, we chose a 2 minute treatment time for mapping phosphorylation changes, assuming that any changes seen at that time are not influenced by changes in transcription. Using statistical analysis across 4 biological replicates, we next identified differential phosphosites (note that these are mapped to protein groups – not necessarily individual proteins, if there are multiple close homologs). We optimized the treatment and sample preparation procedure and measured samples on multiple mass spectrometers, and found the results to be very robust. Two minutes IAA treatment on Arabidopsis roots changed the abundance of 1048 phosphosites (FDR ≤ 0.01), where 666 were more and 382 were less abundant after IAA treatment (Figure 1A). We performed shotgun proteomic MS measurement on equivalent samples (Figure 1B), which suggest that observed changes in phosphosite abundance are not generally caused by altered protein levels. Likewise, the differentially phosphorylated proteins showed little overlap with transcriptionally regulated genes (Figure S1A). We therefore regard these changes as hyper-/hypo-phosphorylation events.

**Figure 1:**
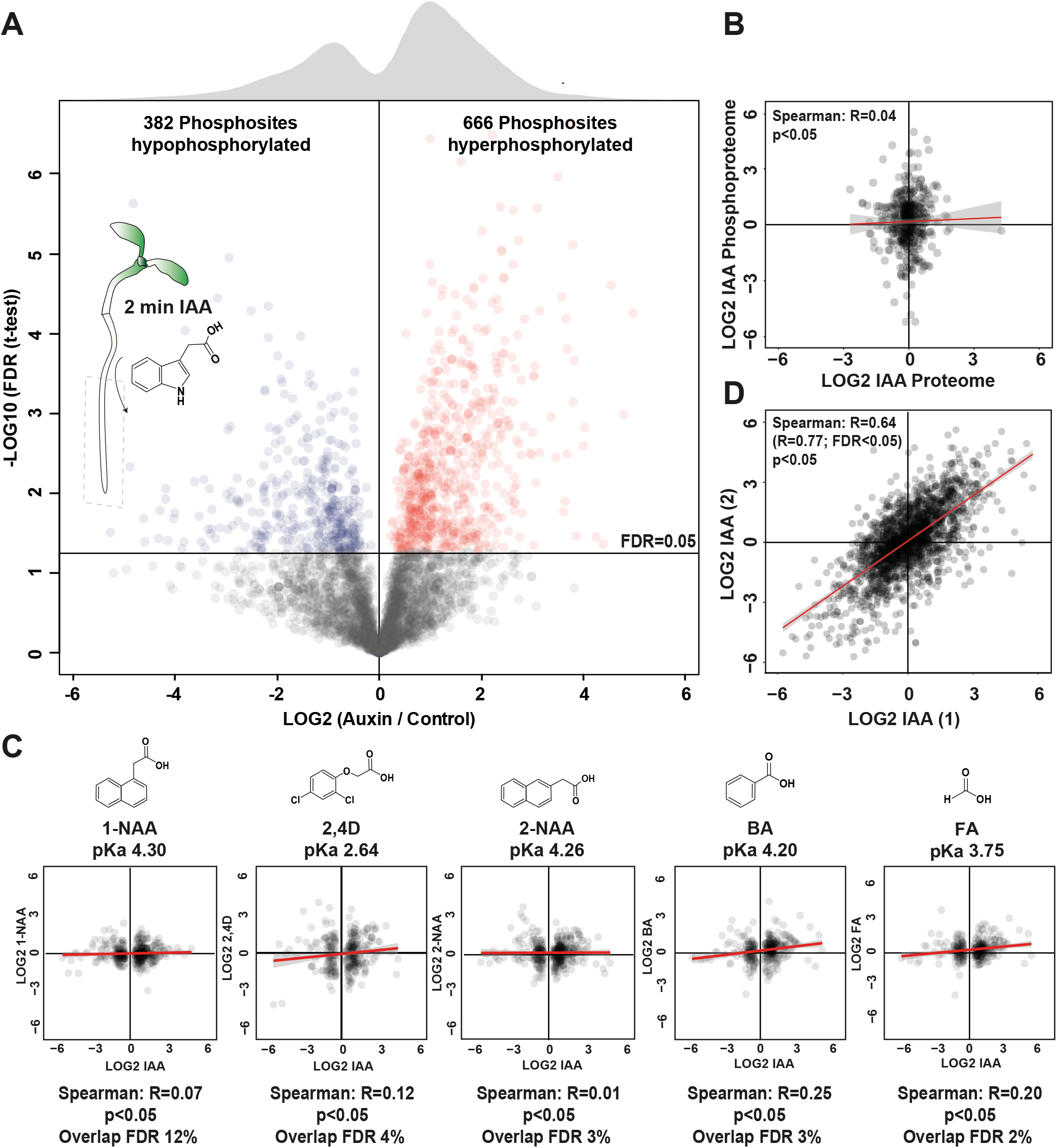
Auxin triggers a unique phosphorylation response. (**A**) Volcano plot showing differential abundance of phosphopeptides between Arabidopsis roots treated with 100 nM IAA, or mock medium for 2 minutes. Each dot represents a single phosphosite, and the significance across replicates is indicated as FDR. Statistical cut-off (FDR≤0.05) is indicated, and a global density histogram is plotted on top. (**B**) Correlation between fold-changes in IAA-trigged shotgun proteome (x-axis) and phosphoproteome (y-axis) both treated with 100 nM IAA for 2 minutes. Red line is regression line with confidence interval (grey). (**C**) Correlation plot of two independent IAA phosphoproteome experiments. Spearman correlation of all sites is 0.64 while it is 0.77 when considering only differential sites at FDR≤0.05. (**D**) Plots comparing differential phosphosites (FDR ≤0.05) in 2 minutes 100 nM IAA treatment (x-axes) with fold-change of corresponding phosphosites in similar treatments with other compounds. Structures and pKa values are given for each compound. Red line indicates regression line (with confidence interval in grey), and Spearman correlation value is indicated in each plot.

The vast effect of a 2-minute IAA treatment on the phosphoproteome to this low IAA concentration is striking, and suggest this to be a hormonal response. However, auxin is a weak organic acid derived from Tryptophan, and it is possible that (part of) the response observed is an unspecific response to weak organic acids or auxin-like molecules. To test chemical specificity, we therefore used a panel of related chemicals (all at 100 nM) in the same set-up, and measured phosphoproteomes after 2 minutes treatment. None of the synthetic auxin analogs 2,4-Dichlorophenoxy-acetic acid (2,4D), 1-Naphtaleneacetic Acid (1-NAA) or 2-Naphtaleneacetic Acid (2-NAA) trigger phosphorylation changes that showed any correlation to those induced by IAA (Figure 1C; Spearman’s rank correlation [R]: 0.12 for 2,4-D; 0.07 for 1-NAA and 0.01 for 2-NAA). Likewise, neither Benzoic Acid (BA; R=0.25) nor Formic Acid (FA; R=0.20) induced IAA-like phosphorylation changes (Figure 1C). As a control, two entirely independent IAA treatments and measurements (Figure 1D) showed strong correlation (R=0.77). Thus, IAA response is chemically specific. Synthetic auxins have auxinic activity in several physiological assays, and it is therefore striking that 2,4-D, 1-NAA and 2-NAA failed to trigger IAA-like phosphorylation changes. To test if these can act like IAA in this response, albeit less efficiently, we measured phosphoproteomes at a 10-fold higher concentration (1 μM). Indeed, at this concentration, 1-NAA (R=0.90) and 2-NAA (R=0.88) could induce IAA-like phosphorylation changes, but BA could not (R=0.12; Figure S1B). Thus, IAA induces a rapid, chemically specific, hormonal phosphorylation response in Arabidopsis roots.

### Auxin phospho-response is ultra-rapid and dynamic

To explore the temporal dynamics of the newly identified phosphorylation response, we generated a time series of IAA treatments on Arabidopsis roots, ranging from 30 seconds to 10 minutes. To ensure that this time series was not confounded by auxin-independent effects of submerging roots in growth medium, we also sampled solvent mock controls for each time point, and subtracted the phosphosite abundance in the mock treatment from each IAA treatment (Figure 2A). This led to a set of unique phosphoproteomes that could be clearly resolved by Principal Component Analysis (PCA; Figure 2B). Strikingly, even at 30 seconds, IAA triggered changes in abundance of hundreds of phosphosites (Figure 2D), underlining the rapid nature of the response. Compiling all differentially phosphorylated sites across the entire time series, a total of 2962 phosphosites is regulated by IAA (Figure 2D), corresponding to 1770 proteins, representing ∼5% of the proteins encoded by the Arabidopsis genome.

**Figure 2:**
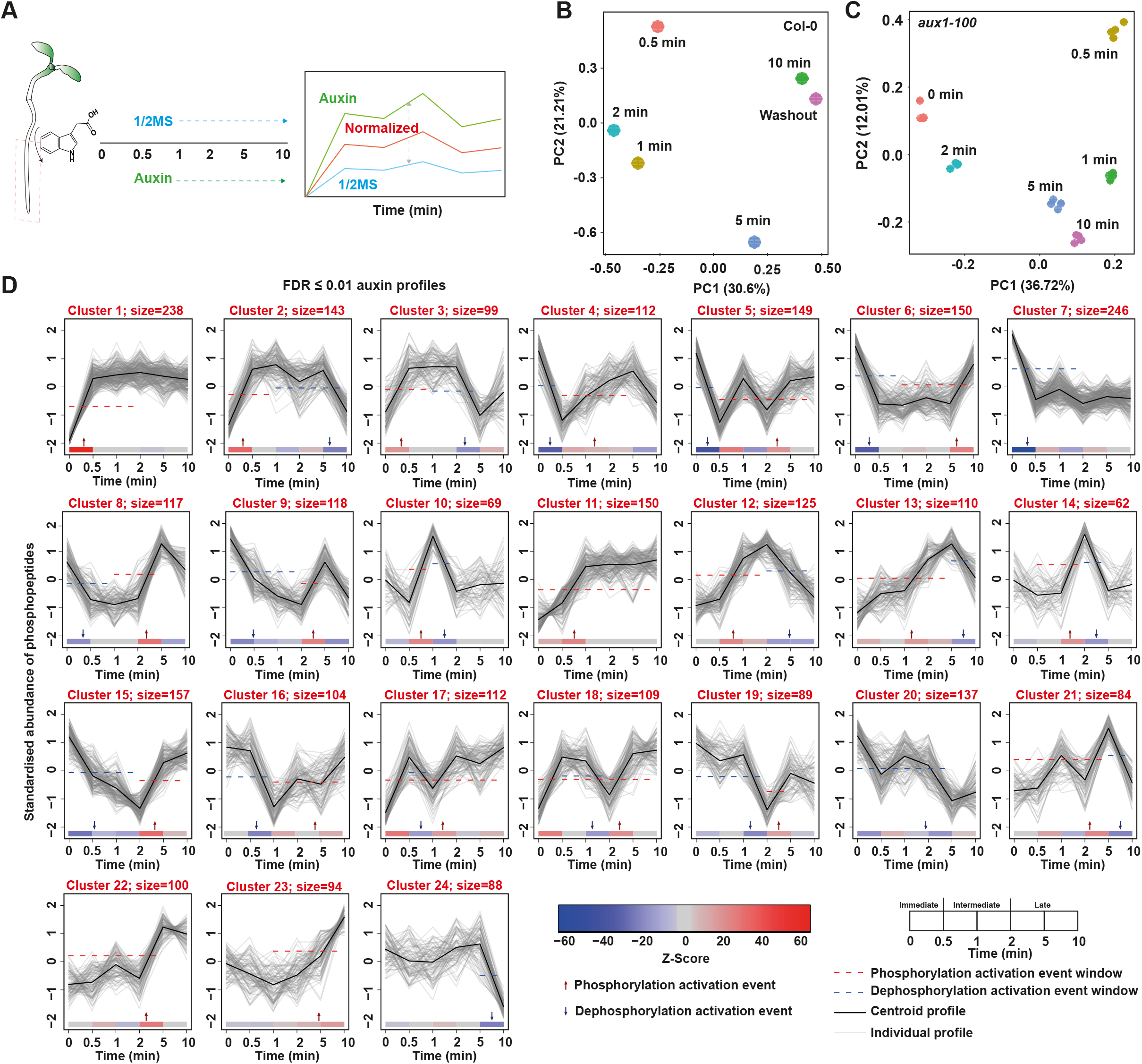
Dynamics of auxin phosphoresponse. (**A**) Schematic overview of treatment, time and analysis procedure. Roots of Arabidopsis seedlings were treated with 100 nM IAA or 1/2MS medium with DMSO as treatment control for various timepoints. Datasets were individually analyzed and intensities of treatment control were subtracted from the auxin responsive profiles (FDR ≥ 0.01) resulting in normalized auxin-responsive profiles. (**B**) Principal component analysis of normalized auxin-responsive profiles and washout. (**C**) Principal component analysis of normalized MS1 phosphosite intensities of time series in the *aux1-100* mutant. (**D**) Time series ordering of all FDR ≥ 0.01 log2 Z-scored normalized auxin-responsive profiles using the Minardo-model. Clusters are ordered based on earliest phosphorylation event. Phosphorylation event is based on the median time at which all individual profiles in a cluster cross half-maximal abundance within each event window (identified by the red or blue dashed line in graph and red and blue arrows on x-axis, respectively).

We next used the Minardo-Model^44^ to order groups of phosphopeptides that show similar trends of phosphorylation (Figure 2D). In each cluster, generalized linear models are derived from individual profiles which, together with Z-scores and post hoc Tukey test, infers event windows at which the majority of profiles show half-maximal amplitude response. From this analysis, the earliest events likely occur well within 30 seconds after treatment (Figure 2D; Clusters 1-9). This analysis clearly identified a range of temporal patterns in IAA-dependent phosphorylation. These ranged from transient, early or late hyper- or hypo-phosphorylation to gradual hyper- or hypo-phosphorylation, and oscillatory hyper- and hypo-phosphorylation (Figure 2D).

Phosphorylation events are often counteracted by regulatory dephosphorylation in enzymatic feedback loops. Given the highly dynamic phosphorylation patterns upon IAA treatment, it is possible that such feedbacks are operating here. To test the reversibility of the widespread IAA-induced differential phosphorylation, we performed a 2-minute IAA treatment, followed by 5-minute washout treatment with IAA-less medium. If the response were reversible, the profile after washout should resemble that after 2 minutes, or even before IAA treatment. Instead, PCA analysis of the washout treatment, along with the time series, grouped the washout treatment with the 10-minute treatment (Figure 2B). This suggests that the IAA response is not globally reversible and persists for minutes even after removal of the signal.

### Genetic requirements of rapid auxin response

The widespread phosphorylation response to IAA treatment is novel, and a key question therefore is what mechanism mediates it. We initially explored the likely location of the receptor or binding site for IAA. As a weak acid, IAA tends to be protonated in the apoplast, and can enter the cell to some degree. Import into the cytosol is however facilitated by the AUX1 permease^45^. To ask if the binding site for IAA is intracellular or apoplastic, we measured a time series phosphoproteome in the *aux1-100* mutant (Figure 2C). In case auxin is perceived intracellularly, one would expect a delay in the response. The *aux1* mutant did show a substantially different phosphorylation pattern, and PCA analysis showed clear separation of wild-type and mutant, irrespective of IAA treatment (Figure 2C). Importantly however, the *aux1* mutant showed a profound response to IAA, even at 30 seconds, which suggests that facilitated IAA influx is not strictly required for fast response. Hence, at least some of the IAA binding capacity for this fast response likely resides in the apoplast.

Two auxin binding sites have been studied in detail: the nuclear TIR1/AFB receptors mediate auxin-dependent gene regulation, as well as some rapid non-transcriptional responses^14,26,28^. For the latter, the cytosolic AFB1 protein appears to play a prominent role^14^. The ABP1 protein resides both in the endoplasmic reticulum and in the apoplast, and has been connected to non-transcriptional responses, likely through interactions with the TMK1 receptor-like kinase^16^. We compared previously recorded^16^ phosphoproteomes in *abp1* and *tmk1* mutants with one generated of the *afb1* mutants after a 2-minute IAA treatment.

Strikingly, each of the three mutants showed profound changes in their auxin-induced phosphoproteomes (Figure 3A,C). In all three mutants, a significant fraction of the auxin-triggered changes in phosphorylation were lost, or in cases, oppositely regulated (Figure 3A). To address that effect of each mutation on the level of phosphorylation of IAA-regulated sites in the absence of external IAA treatment, we compared mock-treated mutants with wild-type and evaluated the phosphorylation level of the entire set of IAA-triggered differential sites. This analysis showed that IAA-induced phosphosites are strongly hypophosphorylated in untreated *abp1* and *tmk1* mutants, but not in *afb1* (Figure 3B). Thus, *abp1* and *tmk1* accumulate phosphorylation defects during their normal growth, while this is not the case for *afb1*. Quantitatively, the number of IAA-triggered differential phosphosites was strongly reduced in all three mutants (Figure 3D). When clustering mutants according to the similarity of the changes in phosphorylation levels across the entire set of IAA-dependent phospopeptides (in wild-type), we found that *abp1* and *tmk1* clustered together, and *afb1* showed a distinct effect on the global IAA-triggered phosphoproteome (Figure 3E). Strikingly, a large set of phosphosites was similarly affected in *abp1* and *tmk1* mutants, and oppositely affected in the *afb1* mutant (Figure 3E). This behavior is clearly illustrated by examples of individual phosphosites a range of proteins (Figure 3F). This demonstrates that ABP1, TMK1 and AFB1 are all required for normal auxin-triggered phosphorylation, and that ABP1 and TMK1 act coordinately, while AFB1 generally opposes ABP1/TMK1 action.

**Figure 3:**
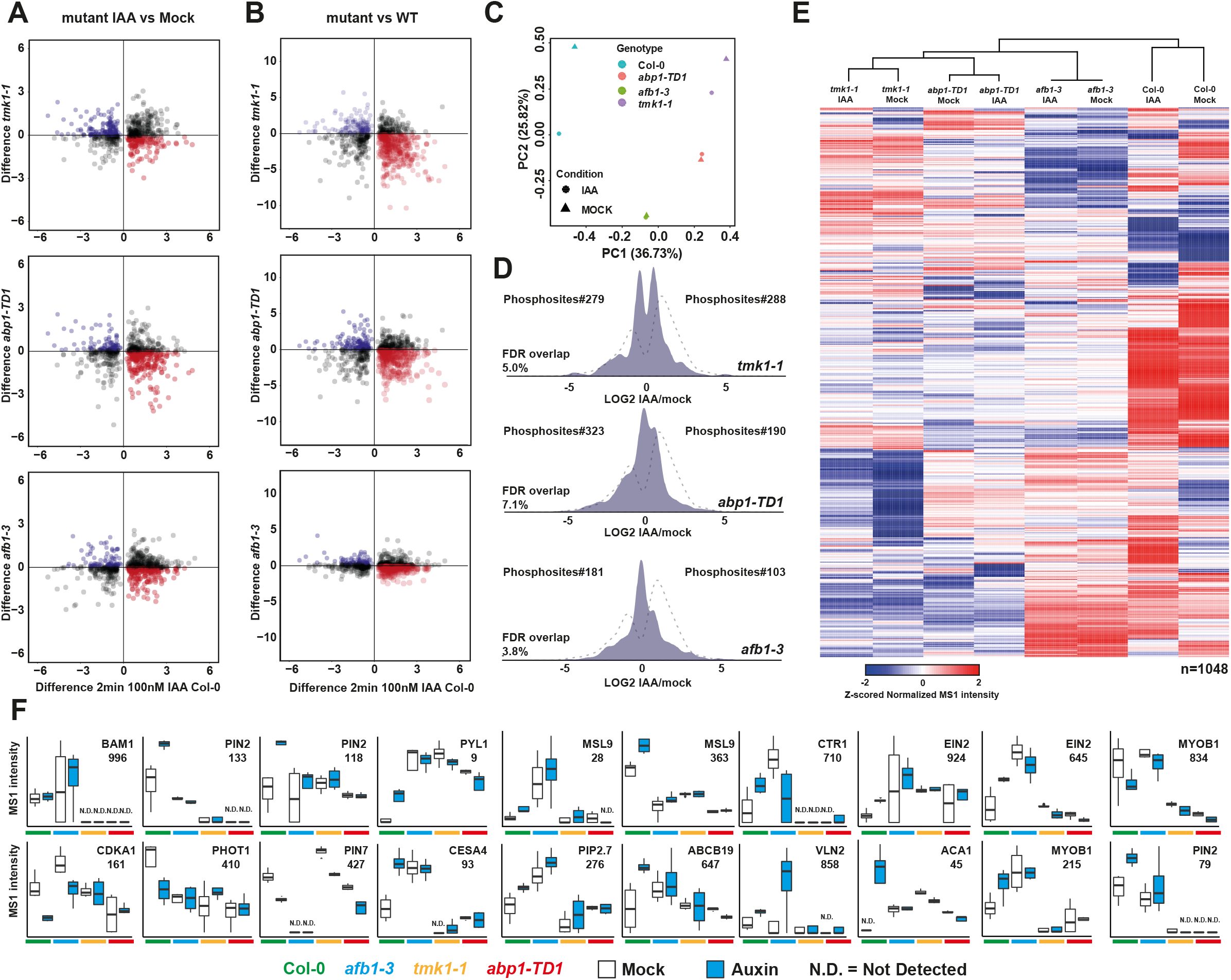
Requirements for auxin-triggered phosphorylation. (**A**) Correlation plot of phosphosite intensity differences between *tmk1-1, abp1-TD1* or *afb1-3* mutants and wild-type after 2 minutes treatments with 100 nM IAA, plotted against fold-change of significantly (FDR ≤0.05) auxin-regulated in Col-0 wild-type. Opposite regulation is marked in red or blue. Similar regulation in black. (**B**) Correlation plot of phosphosite intensity differences between untreated *tmk1-1, abp1-TD1* or *afb1-3* mutants and wild-type (y-axis), plotted against fold-change of significantly (FDR ≤0.05) auxin-regulated in Col-0 wild-type. (**C**) Principal component analysis of mean-normalized MS1 intensities of the 1048 phosphosites that are significantly (FDR ≤0.05) differentially regulated in 2 minutes IAA-treated Col-0, in mock- and IAA-treated wild-type and *abp1, tmk1* and *afb1* mutants. (**D**) Distribution histograms depicting fold-change (IAA/mock) of significantly regulated phosphosites of *tmk1-1, abp1-TD1* and *afb1-3*. Dashed grey line depicts distribution in Col-0 wild-type. (**E**) Heatmap of z-scored normalized MS1 phosphosite intensities in *tmk1-1, abp1-TD1* and *afb1-3* mutants and Col-0 wild-type of the 1048 phosphosites that are significantly (FDR≤0.05) differentially regulated in wild-type upon IAA treatment. (**F**) Boxplots of MS1 intensities of 20 phosphosites in Col-0, *tmk1-1, abp1-TD1, afb1-3* under mock (white) and 2 minutes 100 nM IAA conditions (blue). N.D. means not detected in any of the four biological replicates. Color legend of genotypes is indicated below.

### Auxin-dependent phosphorylation targets multiple cellular pathways

To explore the range of cellular functions potentially targeted by IAA through phosphorylation, we initially mined the full set of differential phosphosites for overrepresented functions, families and pathways. Gene Ontology (GO) analysis of differential phosphosites (Figure 4A) showed that a broad range of cellular functions is subject to regulation. These include a plethora of processes at the plasma membrane or endomembranes, such as endocytosis, vesicle fusion and ion transport, but also prominently featured nuclear processes such as transcription, chromatin structure and splicing. While there is very strong enrichment of plasma membrane, cytosolic and nuclear proteins, most cellular locations are targeted by auxin-triggered phosphorylation. Based on this GO analysis, the prediction is that many aspects of cell function are targeted by IAA-triggered phosphoregulation.

**Figure 4:**
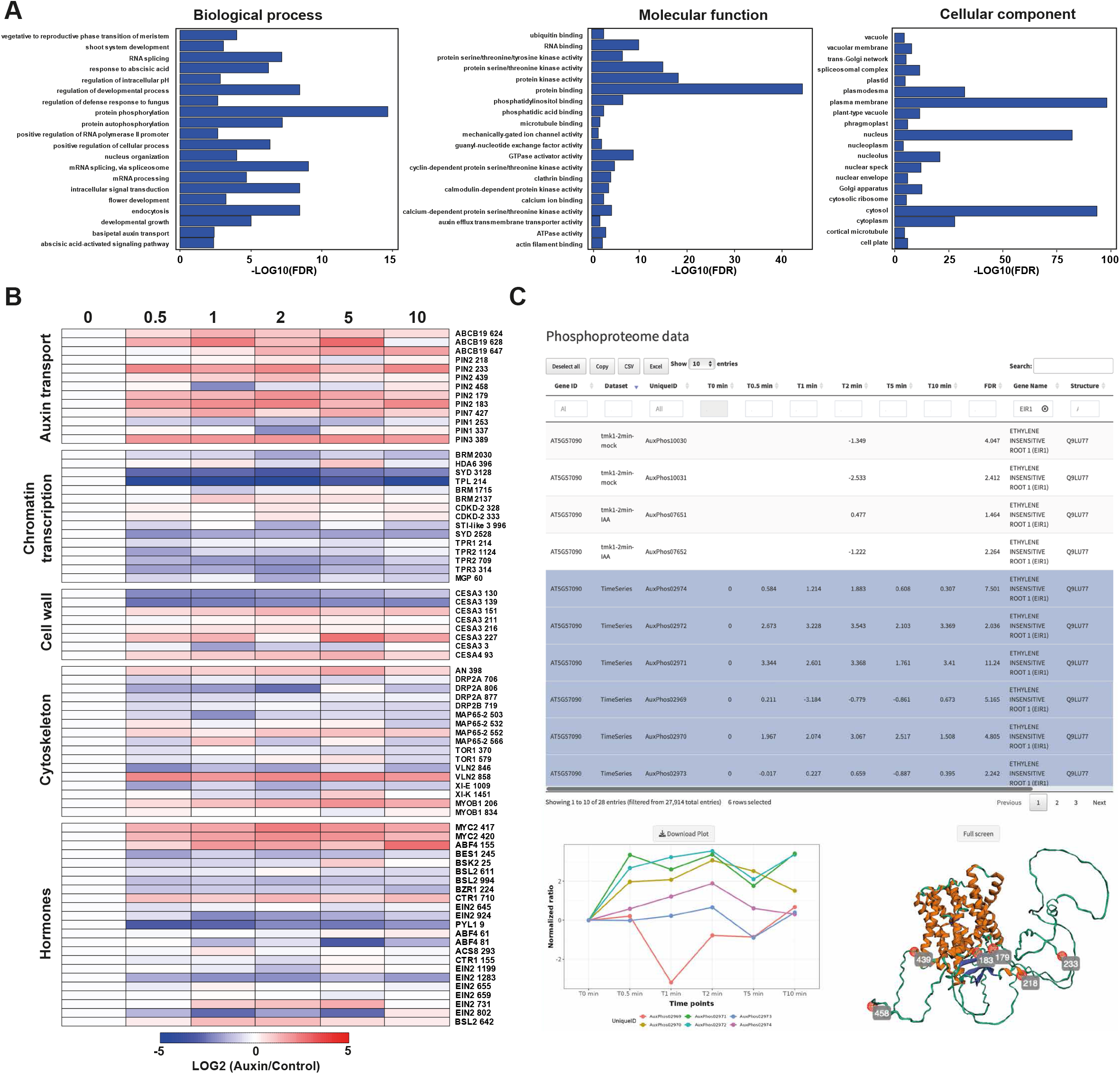
Auxin targets multiple pathways. (**A**) Gene ontology showing the top 20 enrichments of biological process, molecular function and cellular component on the significantly (FDR ≥1.99) auxin-regulated. (**B**) Examples of dynamics of auxin-regulated phosphosites in diverse pathways and functions. Heatmaps depict normalized intensity profiles along the time series. Protein identity is given, along with the position of the phosphosite. (**C**) Screenshots of the R shiny app Auxphos. Selected profiles of a single protein can be selected, visualized in a plot and phosphosites can be mapped on a 3D predicted protein structure using AlphaFold.

We next surveyed the dynamic phosphorylation of proteins within functional groups that had been linked to auxin action, phosphorylation or were clearly enriched in the GO analysis (Figure 4B). PIN phosphorylation is a well-known regulatory step in transporter activation or polar localization. We identified a range of phosphosites on PIN1, 2, 3 and 7. While most were rapidly hyperphosphorylated, those on PIN1 were hypophosphorylated. Interestingly, these all represent sites that have not previously been connected to PIN regulation. We likewise found rapid hyperphosphorylation on the ABCB19 auxin transport protein.

Auxin profoundly regulates gene expression, a function that has thus far been connected to the NAP system^21^. We noticed that there is a very rapid hypophosphorylation on a range of proteins involved in chromatin structure and activity. This includes BRM and SYD SWI/SNF ATPases and nearly all members of the TPL/TPR family. The function of these events is not known, but there it is likely that early phosphorylation events prime or condition later transcriptional responses.

A third functional group that is strongly affected by auxin is cell wall and cytoskeletal proteins. We found many differential phosphosites on Cellulose Synthases, Microtubule Associated Proteins, Myosins, and Dynamin Related Proteins. There was no clear pattern in hyper/hypo-phosphorylation among these, and in some cases, the same protein was oppositely affected at different residues.

Lastly, IAA appears to trigger a range of phosphorylation events in mediators of signaling for other plant hormones. For example, multiple proteins in the ethylene pathway (ACS8, CTR1, EIN2) are targeted, as are multiple brassinosteroid (BES1, BZR1, BSL2, BSK2) and ABA (ABF4, PYL1), and the jasmonic acid component MYC2. Thus, while most of these sites are not yet connected to activity of these factors, it is evident that rapid IAA signaling has the potential to influence the response to other plant hormones.

The phosphorylation changes reported here offer a rich source of data that allows developing hypotheses for future studies. To facilitate the use of this rich dataset and to help interact with the data, we designed a webtool (AuxPhos; https://weijerslab.shinyapps.io/AuxPhos) and user interface. AuxPhos allows to search individual proteins by their unique identifier, and visualize the quantification of its phosphopeptides across the various datasets (wild-type with IAA and various related chemicals; time series; *aux1, abp1, tmk1* and *afb1* mutants). In addition to offering a searchable interface for navigating phosphoproteins, we have implemented AlphaFold2-based protein structural models to visualize on predicted protein structures, where differential phosphorylation occurs. An examples of such visualizations on the EIR1/PIN2 protein are shown in (Figure 4C). As further phosphoproteomic data will become available, these will be integrated in the tool.

### Identification of kinases in auxin-triggered phosphoresponse

Given the availability of a densely sampled time series and the identification of nearly 3000 phosposites, we asked if the dataset could be used to infer phosphorylation relationships. To this end, we filtered the entire phosphoproteome dataset for phosphosites in the (predicted^67^) activation loop of the full set of Arabidopsis kinases (Figure 5A). This identified 26 kinases that were differentially phosphorylated in their activation loop during the IAA-triggered time series (Figure 5B), most of which were hyperphosphorylated, likely signifying activation of kinase activity. We next performed a regression analysis (Figure 5A) where we identified phosphorylation sites that temporally matched or followed the dynamics of activation of each kinase, obeying several statistical criteria. This led to the generation of a kinase-target network encompassing all 23 regulated kinases and 2140 phosphotargets (Figure S3). To test the validity of this inference approach, we asked if protein kinases with known phosphotargets were correctly predicted. This analysis is challenging because of the relatively small number of well-documented kinase-substrate relationships with information about the exact phosphosite. Nonetheless, we found several such pairs to conform to their predicted relationship in our dataset: A RAF kinase was connected to its OST1 target, and this OST1 kinase to a bZIP target. In addition, D6PK was connected to PIN7 (Figure 5C), consistent with PIN phosphorylation by this kinase^46^.

**Figure 5:**
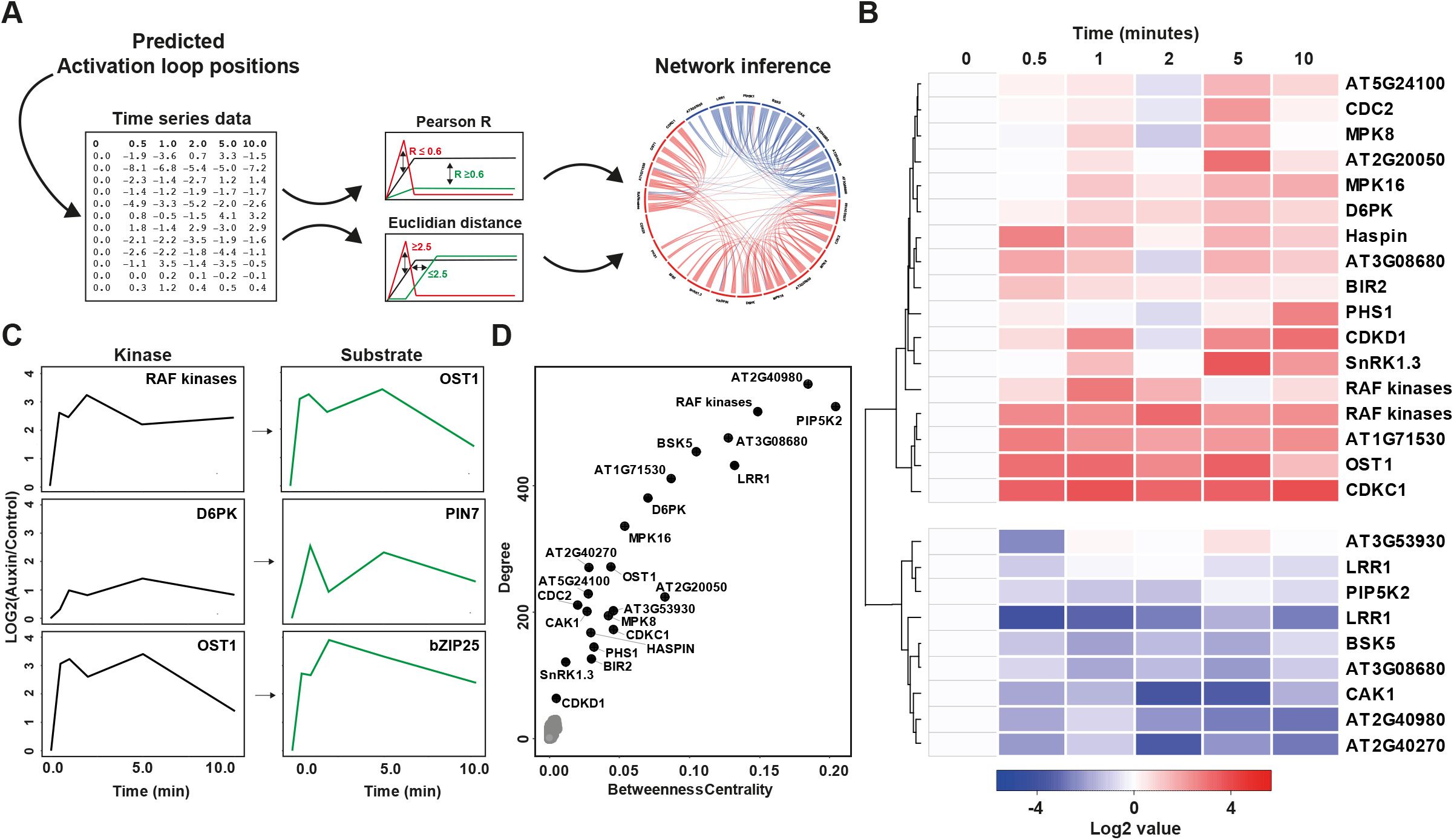
Inference of kinase-substrate relationships. (**A**) Schematic overview of the kinase network inference approach. Phosphosites in predicted and known activation loops were selected from the time series data. The time series profile of the individual activation loop phosphosites were next analysed using correlation analysis (Pearson R and Euclidian distance). Profiles of phosphosites passing the threshold (Pearson R ≥0.6, Euclidian distance ≤2.5) were considered as potential substrates, and used to build a network. (**B**) Heatmap depicting the normalized intensity profiles of phosphopeptides in the activation loop of kinases along the time series. (**C**) Plots showing normalized phosphopeptide abundance profiles along auxin time series of known kinase-substrate pairs recovered in the network inference approach. (**D**) Network position of auxin-regulated kinases analyzed in the inference approach. Plot depicts degree (i.e. how many edges/interactions a node has) and betweenness centrality. The latter is a measurement of hub importance/centrality of a node.

We next ranked the 23 kinases according to their weight and position in the phosphonetwork (Figure 5D), indicated by their degree and their betweenness centrality. These two parameters were correlated in this network. This identified several kinases as potential hubs in this network. These include the D6PK kinase, as well as the LRR-RLK protein LRR1, a group of closely related RAF kinases and PIP5K2. The latter is a phosphoinositide kinase, and it is an open question if the PIP5K activity or the membrane changes it induces are correlated with downstream phosphorylation. In any event, this network analysis offers a prioritized set of auxin-regulated kinases that are strong candidates for mediating rapid responses.

### Rapid auxin response controls membrane polarization

The many functions enriched in the IAA-dependent phosphoproteome include the control of intracellular pH (Figure 4A), which we explored further given that membrane polarity and apoplastic pH have been shown to be subject to auxin control^14,15,27,29,47^. To explore this regulation in more detail, we analyzed phosphorylation dynamics of the family of Arabidopsis H+-ATPase (AHA) proton pumps. These directly transport protons across membranes and tend to be localized to the plasma membrane, thus directly controlling apoplastic pH and membrane polarity^14,27,29^. One AHA is directly phosphorylated by TMK1^27,29^, and we indeed found both TMK1 and AHA1/2 phosphorylation to be rapidly induced (Figure 6A). Likewise, most AHA proteins are differentially phosphorylated within 30 seconds, followed by protein-specific temporal dynamics. This suggests a family-wide regulation of AHA pumps, and predicts that IAA treatment triggers profound and very rapid apoplastic pH changes. Previous pH imaging showed that IAA acts on apoplastic pH within minutes^27^, but did not allow to dissect early kinetics. We therefore used the pH-sensitive HPTS dye^47^ to measure root surface pH changes in a line expressing the cytosolic calcium sensor R-GECO1^48^. We imaged roots in a microfluidic device mounted on a vertical microscope to record the apoplastic pH response in the seconds immediately following auxin application. We found that IAA treatment led to an almost instantaneous apoplastic alkalinization, represented by an increase in HPTS fluorescence excited by 488 nm laser (Figure 6B). After reaching a peak at 90 seconds, the surface pH decreased to a new plateau which was more alkaline than the pre-treatment levels. This behavior is consistent with the hypothesis that auxin causes a proton influx into cells, which is counteracted by the activation of proton extrusion from the cells^27^, and this two-phase process is reflected in the dynamics of AHA phosphorylation. Their complex phosphorylation pattern likely results from a direct activation by the TMKs as well as indirect activation by auxin-induced ion fluxes.

**Figure 6:**
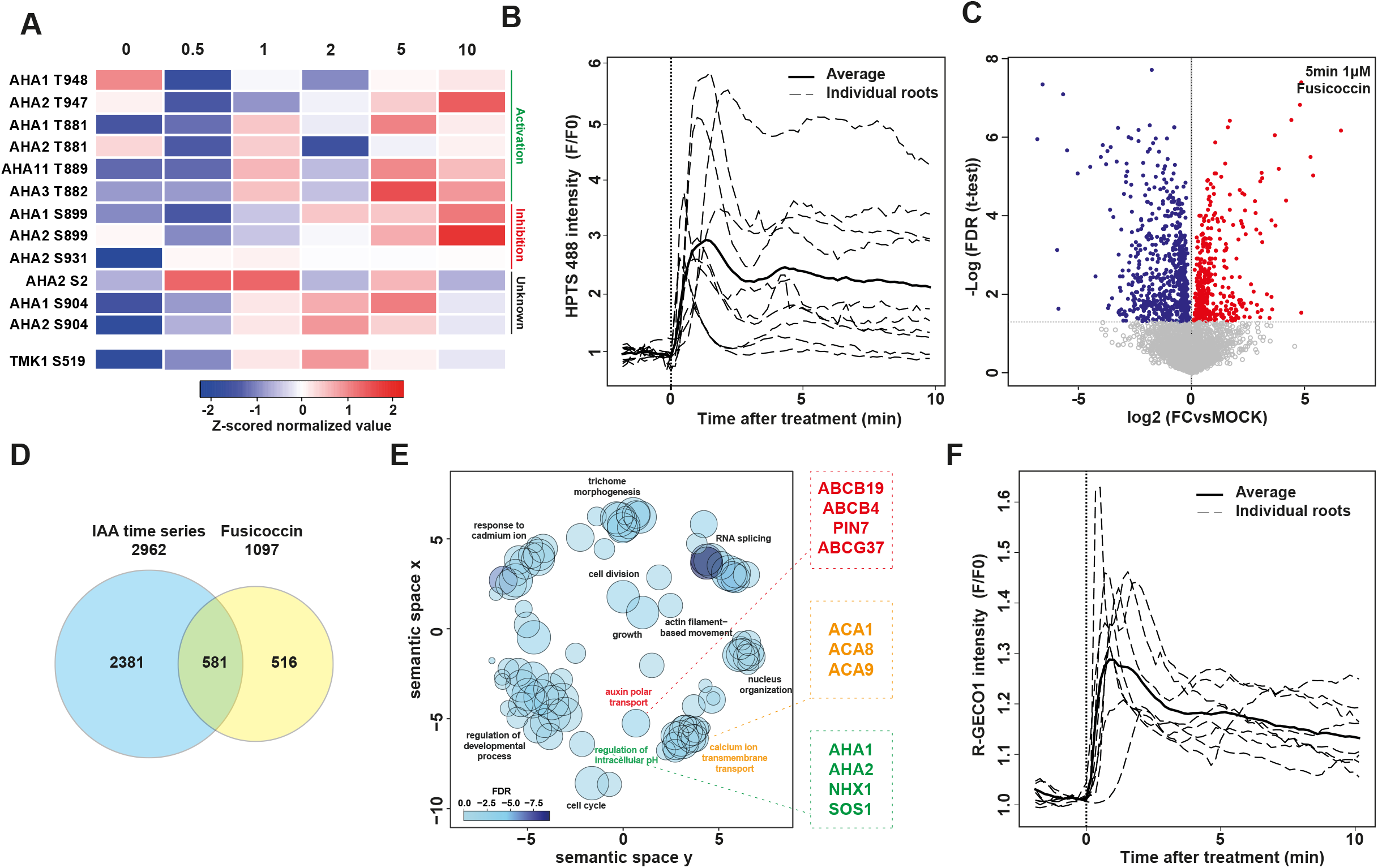
Auxin-triggered apoplastic pH changes. (**A**) Heatmap depicting Z-score normalized values of all identified AHA phosphosites, along the time series, and clustered by identity of the sites. Some represent known activation or inhibition sites, while others are not functionally understood (unknown). A TMK1 phosphosite is shown below for comparison. (**B**) Quantitative analysis of rapid apoplastic pH changes (HPTS fluorescence intensity) following auxin treatment. Dotted lines represent average values from both root sizes of individual roots while black lines represent average of all measured roots (n=6). (**C**) Volcono plot showing phosphosites responsive to 5 minutes 1 μM Fusicoccin (FC) treatment, compared to mock control. (**D**) Venn diagram depicting overlap between significant (FDR ≤0.05) differential phosphosites in the auxin-induction time series, and the FC. (**E**) REVIGO plot (GO analysis) of the 581 phosphosites in the overlap between auxin and FC treatment. Some proteins in highlighted categories are indicated in boxes. (**F**) Quantitative analysis of calcium influx following auxin treatment in the R-GECO1 line. Dotted lines represent normalized values of individual roots while black lines represents average of all measured roots (n=6).

The finding that IAA treatment triggers very rapid pH changes, raises the possibility that the pH changes are themselves causal to some of the later phosphorylation events observed after IAA treatment. To address this question, we treated seedlings with the fungal toxin Fusicoccin (FC), a compound that causes continuous activation of H+-ATPases^50,52^, and recorded phosphoproteomes on dissected roots. A 5-minute FC treatment indeed induced differential phosphorylation on 1097 proteins (Figure 6C). When comparing these phosphosites to those triggered by IAA, we found that more than of half the FC-induced changes are also triggered by IAA, and about 20% of the IAA-induced changes are also triggered by FC (Figure 6D). This suggests that IAA triggers a unique and specific early set of phosphorylation changes, that includes regulatory changes to the H+-ATPases. The changes in pH that ensue from this ATPase regulation then themselves in part trigger a second wave of phosphorylation changes. The FC & IAA-triggered phosphosites include the AHA proteins themselves, suggesting feedback regulation (Figure 6E), but also include PIN and ABCB auxin transporters and ACA Calcium ATPases. Accordingly, IAA treatment also led to near instantaneous calcium influx, which reached a peak after 60 seconds and then reverted to a new plateau (Figure 6F, Supplemental Figure 4), which, similarly to the surface pH, was higher than the pre-treatment level. This is consistent with the tight connection between the regulation of pH and calcium signaling in plant cells^49^. In conclusion, IAA triggers a highly dynamic, and likely feedback-regulated set of changes in membrane ion fluxes and the dynamics of these events is well reflected in the phosphoproteome dataset.

## DISCUSSION

Through phosphoproteomic profiling, we have identified a previously unknown and unsuspected branch of auxin activity. Strikingly, Arabidopsis root cells are capable of changing the phosphorylation state of more than 1700 proteins within 10 minutes of treatment with the natural auxin IAA, many of which respond within 30 seconds. We have not explored yet earlier timepoints, as these become technically impractical to harvest and perhaps effectivity will also be limited by the speed of IAA diffusion into roots. Computational inference suggests that the earliest responses occur around 20 seconds, a timeframe that is consistent with the kinetics of insulin-triggered phosphorylation in animal cells^35^. This auxin-triggered phosphorylation response is remarkably chemically specific, with natural IAA being substantially more effective than synthetic analogs, and related – but physiologically inactive – compounds being unable to trigger the response. This suggests a specific auxin binding site for this response. We explored the role of the cytoplasmic and nuclear AFB1 receptor, as well as the extracellular ABP1 protein, along with its interaction partner TMK1. These analyses show that matters are not simple: all three proteins are required for IAA-dependent phosphorylation, but in different ways. Globally, the *afb1* mutation has essentially opposite effects to either *abp1* or *tmk1* mutants, which are highly similar^16^. Genetically speaking, these could therefore be interpreted as antagonistic components converging on a shared downstream process. At what level these intersect is a question for future investigation, but it is striking that also with regards to controlling apoplastic pH, the nuclear auxin response and TMK1-dependent response have opposite activities^27^.

One caveat of the analysis is that it relies on adding exogenous IAA for observing changes in phosphorylation. One could argue that these treatments do not reflect regulatory events that occur in the absence of external auxin. However, even untreated *abp1* and *tmk1* mutants show hypophosphorylation of many of the sites that are hyperphosphorylated upon IAA treatment in wild-type. This suggests that this same set of sites is controlled by these proteins under normal growth conditions, likely by endogenous IAA, and that defects in their phosphorylation accumulate over time in mutants. The further identification of components in auxin-triggered phosphorylation will likely help to connect these regulatory events to cellular physiology.

The scope of this auxin response is vast, and targets a multitude of proteins, pathways, organelles and functions. Several known or suspected auxin-related proteins are among these, for example PIN and ABCB auxin transporters. Identifying these sites now allows the formulation of specific hypothesis regarding their regulatory role. As a proof of concept in using this resource, we have explored the proton and Calcium fluxes across the plasma membrane that are predicted by the phosphoproteome data. In addition to validating such predictions, we also demonstrate a causal role of apoplastic pH changes in triggering further phosphorylation changes.

A key question is what the impact of this response is on cellular physiology, structure, growth and development. From the many pathways affected, one could expect disruption of the response to have strong phenotypic consequences. It is therefore surprising that the three mutants that strongly affect this response (*abp1, tmk1* and *afb1*) all have largely normal development^16,51,53,54^. Each of these mutants has phenotypic defects in germination (*afb1*)^51^, regeneration (*abp1* and *tmk1*)^16^ or cytoplasmic streaming (*abp1* and *tmk1*)^16^. However, these are subtle. One interpretation is that the phosphorylation network is highly non-linear, with multiple sites being targeted on proteins, phosphatases counteracting kinases, and complex functional redundancies. Given the speed of phosphorylation, it is well possible that the ease of perturbation is balanced by biochemical robustness. It will be interesting to discover the key hubs in the phosphorylation network to understand both the physiological function and the mechanisms underlying robustness. It should be added that this set of mutants has been studied mostly, if not exclusively, under lab-grown, controlled conditions. It is possible that this response system mediates a function that is required in the ecologically relevant setting, and less so in well-controlled conditions.

In relating the phosphorylation response to biology, an important consideration is where this response occurs. We have sampled only roots, for purposes of reproducibility and limiting tissue complexity. It does raise the question of whether similar changes occur – and pathways are affected – in other plant parts. Limitations to sensitivity of mass spectrometry currently make it essentially impossible to capture phosphoproteomes of root cell types, even if just because such analysis require tissue dissociation which would preclude studying rapid responses. When examining the proteins targeted, many are expressed beyond roots, and it would appear reasonable to assume that a response as profound as this is not limited to a single cell type or organ.

When exploring the proteins and functions targeted by IAA-triggered phosphorylation, several stand out. Firstly, mediators of other hormone pathways are prominently regulated. One interesting example is the EIN2 protein^55,56^. This protein is known to be phosphorylated by the CTR1 kinase on two Serines (645 and 924) to control nuclear localization in response to ethylene ^55,56^. Both these sites are hypophosphorylated rapidly following IAA treatment, which predicts that IAA rapidly alters EIN2 localization and ethylene response. Another interesting case is the coordinated hypophosphorylation of a slew of chromatin regulators. This includes BRM and SYD SWI/SNF ATPases and TPL/TPR co-repressors that are all involved in mediating IAA-regulated transcriptional responses^57–59^. The functional impact of these phosphorylations – if any – is unknown. However, it is very likely that the rapid response system interacts with subsequent transcriptional responses, perhaps by priming cells for gene regulation upon prolonged exposure. In any case, our findings reveal the tremendous molecular changes occurring in cells treated with auxin in the minutes prior to the time window during which transcriptional activity is normally recorded.

Lastly, we would like to emphasize the strength of combining ultra-rapid, dense and successive sampling in this approach. We are not aware of other studies using a similar approach to infer the dynamics of any form of rapid signaling in plants. Here, we have leveraged the data to predict kinase-substrate relationships based on regression of temporal profiles. These are successful in capturing known relationships, and help to predict a set of strong candidate kinases that may mediate aspects of auxin-triggered rapid signaling, and will be subject of future investigation. In the accompanying paper (Kuhn, Roosjen et al., accompanying manuscript), we report the identification of a key mediator of rapid phospho-signaling in part through this inference strategy. We expect that similar strategies as reported here will be instrumental in resolving causalities and identifying regulators in responses to other signals, whether hormones, endogenous peptides or environmental or microbial signals.

With reporting this novel, unsuspected and vast response, we offer a new avenue in studying a profoundly important plant signaling molecule. Not only is auxin a major regulator of “normal” plant growth and development, it is also a potent herbicide and growth regulator in agricultural and horticultural practice. We anticipate that mining the rich datasets, compiled in the AuxPhos tool (https://weijerslab.shinyapps.io/AuxPhos) will be the starting point of many studies that help understand the mechanistic basis of the ability of auxin to control plant cell biology, physiology, growth and development.

## ACKNOWLEDGEMENTS

We are grateful to Mark Estelle for making the *afb1* mutant available, our team members for discussions and suggestions. This work was supported by funding from the Netherlands Organization for Scientific Research (NWO): VICI grant 865.14.001 grant to D.W. and VENI grant VI.VENI.212.003 to A.K.; the European Research Council AdG DIRNDL (contract number 833867), StG CELLONGATE (contract 803048) to M.F. and AdG ETAP (contract 742985) to J.F.; and the Austrian Science Fund (FWF, P29988) to J.F..

## AUTHOR CONTRIBUTIONS

Conceptualization: M.R., A.K., J.F., D.W.; Methodology: M.R., S.B.; Software: S.K.M., J.K.; Formal analysis: M.R., P.K.; Investigation: M.R., A.K., P.K.; Writing – Original Draft: M.R., A.K., D.W.; Writing – Review & Editing: all authors; Visualization: M.R., S.K.M., P.K.; Supervision: M.F., J.F., D.W.; Funding Acquisition: A.K., M.F., J.F., D.W..

## MATERIALS AND METHODS

### Plant material, growth and treatments

Arabidopsis seeds were surfaced sterilized, stratified in 0.1% agarose and grown on top of a 100 μm nylon mesh on half-strength Murashige and Skoog (MS) medium with 0.8% Agar (Duchefa). Seedlings were vertically grown in a growth chamber at 22 °C under long days (16h:8h light:dark) for 5 days. For rapid auxin treatment, 100 nM IAA (dissolved in DMSO) or an equal volume of DMSO was diluted in liquid half-strength MS medium. Plates were treated one at a time, placed horizontally, and solution was slowly added until roots were submerged. Plates were subsequently placed vertically again, to minimize gravitropic stimulation. Roots were harvested by scalpel and directly frozen in liquid nitrogen. Roots were ground to a fine powder and kept at -80 °C until further processing. For comparisons between chemicals, timepoints and genotypes, all replicates of all treatments were grown on the same day and processed independently.

Columbia-0 was used as wild-type in all experiments. Mutants used were *abp1-TD* ^60^, *tmk1-1*^54^, *afb1*-*3*^61^ and *aux1*-*100*^62^. For Calcium imaging, a line expressing the R-GECO1 sensor was used^48^.

### Chemicals

For microscopy, IAA (10 mM stock in 96% ethanol; Sigma Aldrich), HPTS (8-Hydroxypyrene-1,3,6-Trisulfonic Acid, Trisodium Salt, 100 mM stock in water; Thermo Fisher Scientific, H348) were used. For phosphoproteomics, IAA (10 mM stock in DMSO; Alfa Aesar), 1-NAA (10 mM stock in DMSO; Sigma Aldrich), 2-NAA (10 mM stock in DMSO; Sigma Aldrich), Benzoic acid (10 mM stock in water; Sigma Aldrich), Formic Acid (10 mM stock in water; Sigma Aldrich) and Fusicoccin (1 mM stock in in 96% ethanol; Sigma Aldrich) were used.

### Phosphopeptide enrichment

For phosphopeptide enrichment, ground Arabidopsis roots powder was suspended in an extraction buffer with 100 mM Tris-HCl pH 8.0, 7 M Urea, 1% Triton-X, 10 mM DTT, 10 U/ml DNase I (Roche), 1 mM MgCl_2_, 1% benzonase (Novagen), 1xphosphoSTOP (Pierce) and 1x cocktail protease inhibitor (Pierce). The suspended lysate was sonicated using a cooled (4°C) waterbath sonicator (Qsonica) using 30 cycles of 30 seconds ON and 30 seconds OFF at 90% amplitude. Lysate was subsequently spun down using a cooled (4°C) tabletop centrifuge at 20.000xg for 30 minutes. After centrifugation supernatant was collected and an extra 1% (v:v) of benzonase was added and incubated for 30 minutes at room temperature. Acrylamide was added to 50 mM and incubated for an extra 30 minutes at room temperature. After alkylation, proteins were precipitated using methanol/chloroform. To the lysate, 4 volumes of methanol, 1 volume of chloroform and 3 volumes of milliQ was added with rigorous vortexing in between. Lysate was centrifuged for 10 minutes at 5000 rpm. After centrifugation, the top layer was discarded and 3 volumes of methanol were added to further precipitate the protein layer by centrifugation for 10 minutes at 5000 rpm. After centrifugation, the supernatant was discarded and protein pellet was air dried. Proteins were next resuspended in 50 mM ammonium bicarbonate (ABC) and sonicated using a cooled (4°C) waterbath sonicator (Qsonica) using 30 cycles of 30 seconds ON and 30 seconds OFF at 90% amplitude. After sonication, protein concentration was measured by Bradford reagent (Biorad). For every biological replicate 500 μg protein was digested with sequencing grade trypsin (1:100 trypsin:protein; Roche) overnight at room temperature. Next, peptides were desalted and concentrated using home-made C18 microcolumns. For peptide desalting and concentrating, disposable 1000 μl pipette tips were fitted with 4 plugs of C18 octadecyl 47 mm Disks 2215 (Empore™) material and 1 mg:10 μg of LiChroprep® RP-18 (Merck) : peptides. Tips were sequentially washed with 100 % methanol, 80 % Acetonitrile (CAN) in 0.1% formic acid and twice equilibrated with 0.1 % formic acid. All chromatographic steps were performed by centrifugation for 4 minutes at 1500xg. After equilibration, peptides were loaded for 20 minutes at 400xg. Bound peptides were washed with 0.1% formic acid and eluted with 80 % ACN in 0.1 % formic acid for 4 minutes at 1500xg. Eluted peptides were suspended in loading buffer (80 % acetonitrile, 5 % tri-fluor acetic acid (TFA)). For phosphopeptide enrichment, MagReSyn® Ti-IMAC beads (Resyn bioscience) were used. For every reaction, a 1:4 peptide:bead ratio was used. Beads were equilibrated in loading buffer, resuspended peptides were added and incubated for 20 minutes at room temperature with slow mixing. After 20 minutes, bead-bound phosphopeptides were washed once in loading buffer, once in 80 % acetonitrile, 1 % TFA, and once in 10 % acetonitrile, 0.2 % TFA. After washing, phosphopeptides were eluted twice with x ul 1 % NH_4_OH. After the last elution, phosphopeptides were acidified using 10 % formic acid. Phosphopeptides were subsequently concentrated using home-made C18 microcolumns. For peptide desalting and concentrating, disposable 200 μl pipette tips were fitted with 2 plugs of C18 octadecyl 47 mm Disks 2215 *(Empore™)* material and 1mg:10 μg of LiChroprep® RP-18 (Merck) : peptides. Tips were sequentially washed and equilibrated as described above. After equilibration, peptides were loaded for 20 minutes at 400xg. Bound peptides were washed with 0.1 % formic acid and eluted with 80 % ACN in 0.1 % formic acid for 4 min at 1500xg. Eluted peptides were subsequently concentrated using a vacuum concentrator for 30-60 minutes at 45°C and resuspended in 15μl of 0.1 % formic acid.

### Filter aided sample preparation and peptide fractionation

For FASP 30kDa cut-off amicon filter units (Merck Millipore) were used. Filters were first washed by appling 50μl urea buffer UT buffer (8M Urea and 100mM Tris pH8.5) and centrifuging for 10 minutes on 11000 RPM at 20° C. The desired amount of protein sample (100μg) was next mixed with UT buffer until a volume of 200 μl, applied to the filter and centrifuged for 15 minutes on 11000 RPM at 20° C. Filter was washed with UT buffer by centrifugation for 15 minutes on 11000RPM at 20° C. Retained proteins were alkylated with 50mM acrylamide (Sigma) in UT buffer for 30 minutes at 20°C while gently shaking. Filter was centrifuged and after that washed trice with UT buffer for 15 minutes on 11000RPM at 20°C. Next filter was washed trice in 50mM ABC buffer. After last wash proteins were cleaved by adding sequencing grade trypsin (Roche) in a 1:100 trypsin:protein ratio. Digestion was completed overnight. The following day the filter was placed into a new tube and peptides were eluted by centrifuging for 15 minutes on 11000RPM at 20°C. Further elution was completed by adding two times 50mM ABC buffer and centrifuging for 10 minutes on 11000RPM at 20°C.

FASP digested peptides (10 μg) were submitted to offline in stage-tip high pH reversed phase (Hp-RP) fractionation. For Hp-RP tips, 2 plugs of C18 octadecyl 47mm Disks 2215 (Empore™) material and 1mg:10 *μg* of LiChroprep® RP-18 (Merck) : peptide were added to a 200 μl tip. Tips were washed with methanol for 4 minutes at 1000xg. Next buffer containing 0.1% formic acid and 80% acetonitrile was added and centrifuged for 4 minutes at 1000xg. Final equilibration was achieved with two washes of 0.1% formic acid and two washes of 20mM ammonium formate (Optima®) pH10 for 4 minutes at 1000xg. Peptides were suspended in 20mM ammonium formate pH10 before loading onto Hp-RP tip. Sample was loaded by centrifugation for 20 minutes at 400xg. Peptides were subsequently eluted with ammonium formate buffers containing 5%,8%,11%,18% and 40% ACN.

### Mass spectrometry

For nano liquid chromatography–tandem mass spectrometry (LC–MS/MS) analysis 5 ul of peptide samples were loaded directly onto a 0.10 * 250 mm ReproSil-Pur 120 C18-AQ 1.9 μm beads analytical column (prepared in-house) at a constant pressure of 825 bar (flow rate of circa 700 nL/min) with 1 ml/l HCOOH in water and eluted at a flow of 0.5 ul/min with a 50 min linear gradient from 9% to 34% acetonitril in water with 1 ml/l formic acid with a Thermo EASY nanoLC1000. An electrospray potential of 3.5 kV was applied directly to the eluent via a stainless steel needle fitted into the waste line of a micro cross that was connected between the nLC and the analytical column. Full scan positive mode FTMS spectra were measured between m/z 380 and 1400 on a Exploris 480 (Thermo electron, San Jose, CA, USA) in the Orbitrap at resolution (60000). MS and MSMS AGC targets were set to 300%, 100% respectively or maximum ion injection times of 50 ms (MS) and 30 ms (MSMS) were used. HCD fragmented (Isolation width 1.2 m/z, 28% normalized collision energy) MSMS scans of 2-5+ charged peaks in the MS scan were recorded in data dependent mode in a cycle time of 1.1 s (Resolution 15000, threshold 2e4, 15 s exclusion duration for the selected m/z +/- 10 ppm).

The MaxQuant quantitative proteomics software package was used to analyse LC–MS data with all MS/MS spectra. The following settings were used: peptide and protein FDRC≤C0.01; the proteome of *A. thaliana* (UniProt ID UP000006548) was used as the protein database; maximum missed cleavage was set at 2; variable modifications Oxidation (M), Acetyl (protein N-term), Deamidation (NQ), pPhospho (STY); fixed modification AcrylAmide (C); match between runs and label-free quantification options were selected.

### Data analysis

Maxquant output was analyzed using Perseus or R. For time series analysis, the Maxquant output PhosphoSTY tab was imported in Perseus ^63^. Data was filtered for reverse, contaminants, only identified by site and localization probability of ≥0.75. Intensity values were log2 transformed and filtered to contain at least 75% valid values in one group. Values were subsequently normalized by median column subtraction. Remaining missing values were imputed from a normal distribution using standard settings in Perseus (width: 0.3, down shift: 1.8) (Figure S2A,B). An FDR permutation based ANOVA test was performed to identify significantly changing phosphosite profiles (FDR ≤0.01). To adjust for treatment response, all log2 transformed profiles from mock treatments were merged with the auxin responsive profiles. Mock values were subsequently subtracted from auxin responsive profiles to obtain normalized auxin responsive phosphosite profiles.

Next, temporal ordering/cluster identification of the phosphosite profiles were done using the Minardo-Model in R^44^. Cluster number was determined in a way that most profiles followed the cluster centroid, resulting in 24 clusters.

Gene ontology enrichment was performed using the database for annotation, visualization and integrated discovery (DAVID)^64,65^. For this, UniProt accession codes were used with duplicates removed. As a background, the full *Arabidopsis thaliana* proteome was used. Next REVIGO^66^ and R were used to reduce overlapping GO-terms.

For kinase network analysis, log2-transformed phosphosite profiles of kinases with phosphosites in the activation loop (as described in^67^) were compared against all FDR significant (FDR≤0.01) profiles using Pearson correlation and Euclidean distance (to also include time offset profiles). Profiles passing a Pearson correlation threshold of ≥0.6 and Euclidian distance threshold of ≤2.5 were extracted for further network analysis. For network analysis, UniProt accession codes were taken as an input for Cytoscape. Network analysis was performed in Cytoscape using standard settings. The degree and betweenness centrality were used to determine signaling hub importance.

Adobe illustrator and R, using standard packages, were used for visualization.

### R shiny app

All the phosphosites and the corresponding enrichment data have been imported into the R environment as CSV files. DataTables, reshape2 and dplyr packages were used for data visualization and data wrangling. 3D protein structures of *Arabidopsis thaliana* proteome predicted through the AlphaFold2 program were downloaded from the AlphaFold database hosted at EBI (https://alphafold.ebi.ac.uk/download). These structures were rendered and visualized using the r3dmol package while the plots were generated using ggplot2 package in R.

### Microfluidics experiments

Four-day-old seedlings were placed into a closable microfluidic chip equipped with hydraulic valves and closed by a microscopy cover glass^14^. One channel contained a control solution (non-buffered 1/2 MS, 100 μM HPTS). The second channel contained a treatment solution (non-buffered ½ MS, 100 μM HPTS, 100 nM IAA). Media flow rate was set to 3 μl/min (OBI1, MFS2 Elveflow and Elveflow software ESI (v.3.04.1). This system was placed to a vertical microscope stage and left for 15 min plant recovery.

### Microscopy imaging

Imaging was performed using a vertical stage Zeiss Axio Observer 7 coupled to a Yokogawa CSU-W1-T2 spinning disk unit with 50 μm pinholes, equipped with a VS-HOM1000 excitation light homogenizer (Visitron Systems). Images were acquired using VisiView software (Visitron Systems, v.4.4.0.14). We used a Zeiss Plan-Apochromat ×10/0.45 objective. HPTS was excited by a 488 nm laser and R-GECO1 by a 561 nm laser. The signal was detected using a PRIME-95B Back-Illuminated sCMOS camera (1,200 × 1,200 pixels; Photometrics). The seedlings were imaged every 10 seconds for a duration of 10 minutes.

### Image analysis

The microscopy images were analyzed in ImageJ Fiji software^68^. HPTS signal intensity was measured using rectangular selection in an area on the outer surface of the root, individually along both sides of the root elongation zone. The intensity of the background was measured using rectangular selection in areas not affected by root response and subtracted from the root sides values. For analysis, the average of the values from both root sides was used. The signal intensity of R-GECO1 was measured in a rectangular selection inside the root at the root elongation zone. Due to the high heterogeneity of the samples, the datasets were normalized using division by initial fluorescence intensity values (F/F0). As initial value F0, we used the average fluorescence intensity at five time points before treatment.

## Data availability

The mass spectrometry proteomics data, protein lists and intensity values of all samples have been deposited to the ProteomeXchange Consortium via the PRIDE^69^ partner repository with the dataset identifier XXX. Source code and complete data used in AuxPhos is available on GitHub (https://github.com/WeijersLab/AuxPhos), while the webtool is accessible from Shinyapps server (https://weijerslab.shinyapps.io/AuxPhos).

## FIGURE LEGENDS

**Supplementary Figure 1: Specificity of auxin-induced phosphorylation**

(**A**) Venn diagram showing overlap of differentially phosphorylated proteins after 2 minutes IAA treatment and differentially expressed genes after 1 hour of IAA treatment (transcriptome data from Kuhn-Roosjen et al., accompanying manuscript). (**B**) Plots comparing differential phosphosites (FDR ≤0.05) in 2 minutes 100 nM IAA treatment (x-axes) with fold-change of corresponding phosphosites in treatments with 1-NAA, 2-NAA or BA at 1 μM. Red line indicates regression line, and Spearman correlation value is indicated in each plot.

**Supplementary Figure 2: Global phosphoprotein properties**

(**A**) Boxplots showing MS1 phosphopeptide intensity distributions across all samples in the time series dataset, and (**B**) normalized and imputed MS1 intensities showing normal distribution of data. (**C**) Principal component analysis of normalized and imputed MS1 intensities show a clear distinction between treatments and the steady state phosphorylation state emphasizing the importance of using a treatment control.

**Supplementary Figure 3: Inferred kinase-substrate network**

Kinase network of the 23 identified kinases with phosphoregulation in their activation loop. Kinases are depicted in blue while substrates are depicted in yellow. Sizes of hexagons are based on degree.

**Supplemental Figure 4: Effect of control medium treatment on Arabidopsis root surface pH and cytosolic Ca2+ level**

(**A**) Root surface pH was visualized by fluorescence intensity of HPTS 488 nm pH-responsive channel. (**B**) The level of cytoplasmic Ca2+ was reported by the fluorescence intensity of R-GECO1. Treatment is indicated by the vertical dotted line (n=3 roots). (**C**) Fluorescence intensity of HPTS 488 nm channel and R-GECO1 was measured in the rectangular regions near the root surface (HPTS) and in the root elongation zone (R-GECO1), as indicated with boxes.

## REFERENCES

1. Went, F.W. (1928). Wuchsstoff und Wachstum. Recueil des travaux botaniques néerlandais 25, 1–116.

2. Went, F.W., and Thimann, K.V. (1937). Phytohormones (MacMillan Company). 3.

3. Friml, J. (2022). Fourteen Stations of Auxin. Cold Spring Harb Perspect Biol 14, a039859. 10.1101/cshperspect.a039859.

4. Morffy, N., and Strader, L.C. (2020). Old Town Roads: routes of auxin biosynthesis across kingdoms. Curr Opin Plant Biol 55, 21–27. 10.1016/j.pbi.2020.02.002.

5. Friml, J., Wiśniewska, J., Benková, E., Mendgen, K., and Palme, K. (2002). Lateral relocation of auxin efflux regulator PIN3 mediates tropism in Arabidopsis. Nature 415, 806–809. 10.1038/415806a.

6. Tao, Y., Ferrer, J.-L., Ljung, K., Pojer, F., Hong, F., Long, J.A., Li, L., Moreno, J.E., Bowman, M.E., Ivans, L.J., et al. (2008). Rapid Synthesis of Auxin via a New Tryptophan-Dependent Pathway Is Required for Shade Avoidance in Plants. Cell 133, 164–176. 10.1016/j.cell.2008.01.049.

7. Scarpella, E., Marcos, D., Friml, J., and Berleth, T. (2006). Control of leaf vascular patterning by polar auxin transport. Genes Dev 20, 1015–1027. 10.1101/gad.1402406.

8. de Rybel, B., Adibi, M., Breda, A.S., Wendrich, J.R., Smit, M.E., Novák, O., Yamaguchi, N., Yoshida, S., van Isterdael, G., Palovaara, J., et al. (2014). Integration of growth and patterning during vascular tissue formation in Arabidopsis. Science (1979) 345. 10.1126/science.1255215.

9. Sachs, T. (2000). Integrating Cellular and Organismic Aspects of Vascular Differentiation. Plant Cell Physiol 41, 649–656. 10.1093/pcp/41.6.649.

10. Casimiro, I., Marchant, A., Bhalerao, R.P., Beeckman, T., Dhooge, S., Swarup, R., Graham, N., Inzé, D., Sandberg, G., Casero, P.J., et al. (2001). Auxin Transport Promotes Arabidopsis Lateral Root Initiation. Plant Cell 13, 843–852. 10.1105/tpc.13.4.843.

11. Dubrovsky, J.G., Sauer, M., Napsucialy-Mendivil, S., Ivanchenko, M.G., Friml, J., Shishkova, S., Celenza, J., and Benková, E. (2008). Auxin acts as a local morphogenetic trigger to specify lateral root founder cells. Proceedings of the National Academy of Sciences 105, 8790–8794. 10.1073/pnas.0712307105.

12. Bates, G.W., and Goldsmith, M.H.M. (1983). Rapid response of the plasma-membrane potential in oat coleoptiles to auxin and other weak acids. Planta 159, 231–237. 10.1007/BF00397530.

13. Etherton, B. (1970). Effect of Indole-3-acetic Acid on Membrane Potentials of Oat Coleoptile Cells. Plant Physiol 45, 527–528. 10.1104/pp.45.4.527.

14. Serre, N.B.C., Kralík, D., Yun, P., Slouka, Z., Shabala, S., and Fendrych, M. (2021). AFB1 controls rapid auxin signalling through membrane depolarization in Arabidopsis thaliana root. Nat Plants 7, 1229–1238. 10.1038/s41477-021-00969-z.

15. Monshausen, G.B., Miller, N.D., Murphy, A.S., and Gilroy, S. (2011). Dynamics of auxin-dependent Ca2+ and pH signaling in root growth revealed by integrating high-resolution imaging with automated computer vision-based analysis. The Plant Journal 65, 309–318. 10.1111/j.1365-313X.2010.04423.x.

16. Friml, J., Gallei, M., Gelová, Z., Johnson, A., Mazur, E., Monzer, A., Rodriguez, L., Roosjen, M., Verstraeten, I., Živanović, B.D., et al. (2022). ABP1–TMK auxin perception for global phosphorylation and auxin canalization. Nature 609, 575–581. 10.1038/s41586-022-05187-x.

17. Ayling, S., and Clarkson, D. (1996). The Cytoplasmic Streaming Response of Tomato Root Hairs to Auxin; the Role of Calcium. Functional Plant Biology 23, 699. 10.1071/PP9960699.

18. Thimann, K. v. (1938). HORMONES AND THE ANALYSIS OF GROWTH. Plant Physiol 13, 437–449. 10.1104/pp.13.3.437.

19. Reinhardt, D., Pesce, E.-R., Stieger, P., Mandel, T., Baltensperger, K., Bennett, M., Traas, J., Friml, J., and Kuhlemeier, C. (2003). Regulation of phyllotaxis by polar auxin transport. Nature 426, 255–260. 10.1038/nature02081.

20. Heisler, M.G., Ohno, C., Das, P., Sieber, P., Reddy, G. v., Long, J.A., and Meyerowitz, E.M. (2005). Patterns of Auxin Transport and Gene Expression during Primordium Development Revealed by Live Imaging of the Arabidopsis Inflorescence Meristem. Current Biology 15, 1899–1911. 10.1016/j.cub.2005.09.052.

21. Weijers, D., and Wagner, D. (2016). Transcriptional Responses to the Auxin Hormone. Annu Rev Plant Biol 67, 539–574. 10.1146/annurev-arplant-043015-112122.

22. Leyser, H.M.O., Lincoln, C.A., Timpte, C., Lammer, D., Turner, J., and Estelle, M. (1993). Arabidopsis auxin-resistance gene AXR1 encodes a protein related to ubiquitin-activating enzyme E1. Nature 364, 161–164. 10.1038/364161a0.

23. Hobbie, L., and Estelle, M. (1995). The axr4 auxin-resistant mutants of Arabidopsis thaliana define a gene important for root gravitropism and lateral root initiation. The Plant Journal 7, 211–220. 10.1046/j.1365-313X.1995.7020211.x.

24. Leyser, O. (1997). Auxin: Lessons from a mutant weed. Physiol Plant 100, 407–414. https://doi.org/10.1111/j.1399-3054.1997.tb03044.x.

25. Abel, S., and Theologis, A. (1996). Early Genes and Auxin Action. Plant Physiol 111, 9–17. 10.1104/pp.111.1.9.

26. Dindas, J., Scherzer, S., Roelfsema, M.R.G., von Meyer, K., Müller, H.M., Al-Rasheid, K.A.S., Palme, K., Dietrich, P., Becker, D., Bennett, M.J., et al. (2018). AUX1-mediated root hair auxin influx governs SCFTIR1/AFB-type Ca2+ signaling. Nat Commun 9, 1174. 10.1038/s41467-018-03582-5.

27. Li, L., Verstraeten, I., Roosjen, M., Takahashi, K., Rodriguez, L., Merrin, J., Chen, J., Shabala, L., Smet, W., Ren, H., et al. (2021). Cell surface and intracellular auxin signalling for H+ fluxes in root growth. Nature 599, 273–277. 10.1038/s41586-021-04037-6.

28. Fendrych, M., Akhmanova, M., Merrin, J., Glanc, M., Hagihara, S., Takahashi, K., Uchida, N., Torii, K.U., and Friml, J. (2018). Rapid and reversible root growth inhibition by TIR1 auxin signalling. Nat Plants 4, 453–459. 10.1038/s41477-018-0190-1.

29. Lin, W., Zhou, X., Tang, W., Takahashi, K., Pan, X., Dai, J., Ren, H., Zhu, X., Pan, S., Zheng, H., et al. (2021). TMK-based cell-surface auxin signalling activates cell-wall acidification. Nature 599, 278–282. 10.1038/s41586-021-03976-4.

30. Narasimhan, M., Gallei, M., Tan, S., Johnson, A., Verstraeten, I., Li, L., Rodriguez, L., Han, H., Himschoot, E., Wang, R., et al. (2021). Systematic analysis of specific and nonspecific auxin effects on endocytosis and trafficking. Plant Physiol 186, 1122–1142. 10.1093/plphys/kiab134.

31. Jin, J., and Pawson, T. (2012). Modular evolution of phosphorylation-based signalling systems. Philosophical Transactions of the Royal Society B: Biological Sciences 367, 2540–2555. 10.1098/rstb.2012.0106.

32. Nurnberger, T., Brunner, F., Kemmerling, B., and Piater, L. (2004). Innate immunity in plants and animals: striking similarities and obvious differences. Immunol Rev 198, 249–266. 10.1111/j.0105-2896.2004.0119.x.

33. Couto, D., and Zipfel, C. (2016). Regulation of pattern recognition receptor signalling in plants. Nat Rev Immunol 16, 537–552. 10.1038/nri.2016.77.

34. Oyama, M., Kozuka-Hata, H., Tasaki, S., Semba, K., Hattori, S., Sugano, S., Inoue, J., and Yamamoto, T. (2009). Temporal Perturbation of Tyrosine Phosphoproteome Dynamics Reveals the System-wide Regulatory Networks. Molecular & Cellular Proteomics 8, 226–231. 10.1074/mcp.M800186-MCP200.

35. Humphrey, S.J., Azimifar, S.B., and Mann, M. (2015). High-throughput phosphoproteomics reveals in vivo insulin signaling dynamics. Nat Biotechnol 33, 990–995. 10.1038/nbt.3327.

36. Kinoshita, T., Caño-Delgado, A., Seto, H., Hiranuma, S., Fujioka, S., Yoshida, S., and Chory, J. (2005). Binding of brassinosteroids to the extracellular domain of plant receptor kinase BRI1. Nature 433, 167–171. 10.1038/nature03227.

37. Wang, X., Kota, U., He, K., Blackburn, K., Li, J., Goshe, M.B., Huber, S.C., and Clouse, S.D. (2008). Sequential Transphosphorylation of the BRI1/BAK1 Receptor Kinase Complex Impacts Early Events in Brassinosteroid Signaling. Dev Cell 15, 220–235. 10.1016/j.devcel.2008.06.011.

38. Ryu, H., Kim, K., Cho, H., Park, J., Choe, S., and Hwang, I. (2007). Nucleocytoplasmic Shuttling of BZR1 Mediated by Phosphorylation Is Essential in Arabidopsis Brassinosteroid Signaling. Plant Cell 19, 2749–2762. 10.1105/tpc.107.053728.

39. Tan, S., Luschnig, C., and Friml, J. (2021). Pho-view of Auxin: Reversible Protein Phosphorylation in Auxin Biosynthesis, Transport and Signaling. Mol Plant 14, 151–165. 10.1016/j.molp.2020.11.004.

40. Zhang, H., Zhou, H., Berke, L., Heck, A.J.R., Mohammed, S., Scheres, B., and Menke, F.L.H. (2013). Quantitative Phosphoproteomics after Auxin-stimulated Lateral Root Induction Identifies an SNX1 Protein Phosphorylation Site Required for Growth. Molecular & Cellular Proteomics 12, 1158–1169. 10.1074/mcp.M112.021220.

41. Nikonorova, N., Murphy, E., Fonseca de Lima, C.F., Zhu, S., van de Cotte, B., Vu, L.D., Balcerowicz, D., Li, L., Kong, X., de Rop, G., et al. (2021). The Arabidopsis Root Tip (Phospho)Proteomes at Growth-Promoting versus Growth-Repressing Conditions Reveal Novel Root Growth Regulators. Cells 10, 1665. 10.3390/cells10071665.

42. Singh, H., Singh, Z., Zhu, T., Xu, X., Waghmode, B., Garg, T., Yadav, S., Sircar, D., de Smet, I., and Yadav, S.R. (2021). Auxin-Responsive (Phospho)proteome Analysis Reveals Key Biological Processes and Signaling Associated with Shoot-Borne Crown Root Development in Rice. Plant Cell Physiol. 10.1093/pcp/pcab155.

43. McClure, B.A., Hagen, G., Brown, C.S., Gee, M.A., and Guilfoyle, T.J. (1989). Transcription, organization, and sequence of an auxin-regulated gene cluster in soybean. Plant Cell 1, 229–239. 10.1105/tpc.1.2.229.

44. Kaur, S., Peters, T.J., Yang, P., Luu, L.D.W., Vuong, J., Krycer, J.R., and O’Donoghue, S.I. (2020). Temporal ordering of omics and multiomic events inferred from time-series data. NPJ Syst Biol Appl 6, 22. 10.1038/s41540-020-0141-0.

45. Yang, Y., Hammes, U.Z., Taylor, C.G., Schachtman, D.P., and Nielsen, E. (2006). High-Affinity Auxin Transport by the AUX1 Influx Carrier Protein. Current Biology 16, 1123–1127. 10.1016/j.cub.2006.04.029.

46. Weller, B., Zourelidou, M., Frank, L., Barbosa, I.C.R., Fastner, A., Richter, S., Jürgens, G., Hammes, U.Z., and Schwechheimer, C. (2017). Dynamic PIN-FORMED auxin efflux carrier phosphorylation at the plasma membrane controls auxin efflux-dependent growth. Proceedings of the National Academy of Sciences 114. 10.1073/pnas.1614380114.

47. Barbez, E., Dünser, K., Gaidora, A., Lendl, T., and Busch, W. (2017). Auxin steers root cell expansion via apoplastic pH regulation in Arabidopsis thaliana. Proceedings of the National Academy of Sciences 114. 10.1073/pnas.1613499114.

48. Keinath, N.F., Waadt, R., Brugman, R., Schroeder, J.I., Grossmann, G., Schumacher, K., and Krebs, M. (2015). Live Cell Imaging with R-GECO1 Sheds Light on flg22- and Chitin-Induced Transient [Ca(2+)]cyt Patterns in Arabidopsis. Mol Plant 8, 1188–1200. 10.1016/j.molp.2015.05.006.

49. Behera, S., Zhaolong, X., Luoni, L., Bonza, M.C., Doccula, F.G., de Michelis, M.I., Morris, R.J., Schwarzländer, M., and Costa, A. (2018). Cellular Ca2+ Signals Generate Defined pH Signatures in Plants. Plant Cell 30, 2704–2719. 10.1105/tpc.18.00655.

50. Olsson, A., Svennelid, F., Ek, B., Sommarin, M., and Larsson, C. (1998). A Phosphothreonine Residue at the C-Terminal End of the Plasma Membrane H+-ATPase Is Protected by Fusicoccin-Induced 14–3–3 Binding. Plant Physiol 118, 551–555. 10.1104/pp.118.2.551.

51. Wang, Y., Goertz, N.J., Rillo, E., and Yang, M. (2022). Negative regulation of seed germination by maternal AFB1 and AFB5 in Arabidopsis. Biosci Rep 42. 10.1042/BSR20221504.

52. Svennelid, F., Olsson, A., Piotrowski, M., Rosenquist, M., Ottman, C., Larsson, C., Oecking, C., and Sommarin, M. (1999). Phosphorylation of Thr-948 at the C Terminus of the Plasma Membrane H ^+^-ATPase Creates a Binding Site for the Regulatory 14-3-3 Protein. Plant Cell 11, 2379–2391. 10.1105/tpc.11.12.2379.

53. Gelová, Z., Gallei, M., Pernisová, M., Brunoud, G., Zhang, X., Glanc, M., Li, L., Michalko, J., Pavlovičová, Z., Verstraeten, I., et al. (2021). Developmental roles of Auxin Binding Protein 1 in Arabidopsis thaliana. Plant Science 303, 110750. 10.1016/j.plantsci.2020.110750.

54. Dai, N., Wang, W., Patterson, S.E., and Bleecker, A.B. (2013). The TMK Subfamily of Receptor-Like Kinases in Arabidopsis Display an Essential Role in Growth and a Reduced Sensitivity to Auxin. PLoS One 8, e60990..

55. Chen, R., Binder, B.M., Garrett, W.M., Tucker, M.L., Chang, C., and Cooper, B. (2011). Proteomic responses in Arabidopsis thaliana seedlings treated with ethylene. Mol Biosyst 7, 2637. 10.1039/c1mb05159h.

56. Ju, C., Yoon, G.M., Shemansky, J.M., Lin, D.Y., Ying, Z.I., Chang, J., Garrett, W.M., Kessenbrock, M., Groth, G., Tucker, M.L., et al. (2012). CTR1 phosphorylates the central regulator EIN2 to control ethylene hormone signaling from the ER membrane to the nucleus in Arabidopsis. Proceedings of the National Academy of Sciences 109, 19486–19491. 10.1073/pnas.1214848109.

57. Kuhn, A., Ramans Harborough, S., McLaughlin, H.M., Natarajan, B., Verstraeten, I., Friml, J., Kepinski, S., and Østergaard, L. (2020). Direct ETTIN-auxin interaction controls chromatin states in gynoecium development. Elife 9. 10.7554/eLife.51787.

58. Szemenyei, H., Hannon, M., and Long, J.A. (2008). TOPLESS Mediates Auxin-Dependent Transcriptional Repression During Arabidopsis Embryogenesis. Science (1979) 319, 1384–1386. 10.1126/science.1151461.

59. Wu, M.-F., Yamaguchi, N., Xiao, J., Bargmann, B., Estelle, M., Sang, Y., and Wagner, D. (2015). Auxin-regulated chromatin switch directs acquisition of flower primordium founder fate. Elife 4. 10.7554/eLife.09269.

60. Gao, Y., Zhang, Y., Zhang, D., Dai, X., Estelle, M., and Zhao, Y. (2015). Auxin binding protein 1 (ABP1) is not required for either auxin signaling or Arabidopsis development. Proceedings of the National Academy of Sciences 112, 2275–2280. 10.1073/pnas.1500365112.

61. Savaldi-Goldstein, S., Baiga, T.J., Pojer, F., Dabi, T., Butterfield, C., Parry, G., Santner, A., Dharmasiri, N., Tao, Y., Estelle, M., et al. (2008). New auxin analogs with growth-promoting effects in intact plants reveal a chemical strategy to improve hormone delivery. Proceedings of the National Academy of Sciences 105, 15190–15195. 10.1073/pnas.0806324105.

62. Bennett, M.J., Marchant, A., Green, H.G., May, S.T., Ward, S.P., Millner, P.A., Walker, A.R., Schulz, B., and Feldmann, K.A. (1996). Arabidopsis AUX1 Gene: A Permease-Like Regulator of Root Gravitropism. Science (1979) 273, 948–950. 10.1126/science.273.5277.948.

63. Tyanova, S., Temu, T., Sinitcyn, P., Carlson, A., Hein, M.Y., Geiger, T., Mann, M., and Cox, J. (2016). The Perseus computational platform for comprehensive analysis of (prote)omics data. Nat Methods 13, 731–740. 10.1038/nmeth.3901.

64. Sherman, B.T., Hao, M., Qiu, J., Jiao, X., Baseler, M.W., Lane, H.C., Imamichi, T., and Chang, W. (2022). DAVID: a web server for functional enrichment analysis and functional annotation of gene lists (2021 update). Nucleic Acids Res 50, W216–W221. 10.1093/nar/gkac194.

65. Huang, D.W., Sherman, B.T., and Lempicki, R.A. (2009). Systematic and integrative analysis of large gene lists using DAVID bioinformatics resources. Nat Protoc 4, 44–57. 10.1038/nprot.2008.211.

66. Supek, F., Bošnjak, M., Škunca, N., and Šmuc, T. (2011). REVIGO Summarizes and Visualizes Long Lists of Gene Ontology Terms. PLoS One 6, e21800. 10.1371/journal.pone.0021800.

67. Montes, C., Wang, P., Liao, C., Nolan, T.M., Song, G., Clark, N.M., Elmore, J.M., Guo, H., Bassham, D.C., Yin, Y., et al. (2022). Integration of multi-omics data reveals interplay between brassinosteroid and Target of Rapamycin Complex signaling in Arabidopsis. New Phytologist 236, 893–910. 10.1111/nph.18404.

68. Schindelin, J., Arganda-Carreras, I., Frise, E., Kaynig, V., Longair, M., Pietzsch, T., Preibisch, S., Rueden, C., Saalfeld, S., Schmid, B., et al. (2012). Fiji: an open-source platform for biological-image analysis. Nat Methods 9, 676–682. 10.1038/nmeth.2019.

69. Perez-Riverol, Y., Bai, J., Bandla, C., García-Seisdedos, D., Hewapathirana, S., Kamatchinathan, S., Kundu, D.J., Prakash, A., Frericks-Zipper, A., Eisenacher, M., et al. (2022). The PRIDE database resources in 2022: a hub for mass spectrometry-based proteomics evidences. Nucleic Acids Res 50, D543–D552. 10.1093/nar/gkab1038.

