## Supplementary figures and images for "An ultra-fast, proteome-wide response to the plant hormone auxin"

### Figure S1

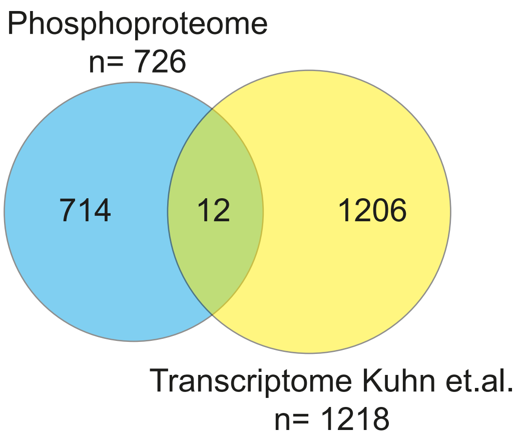

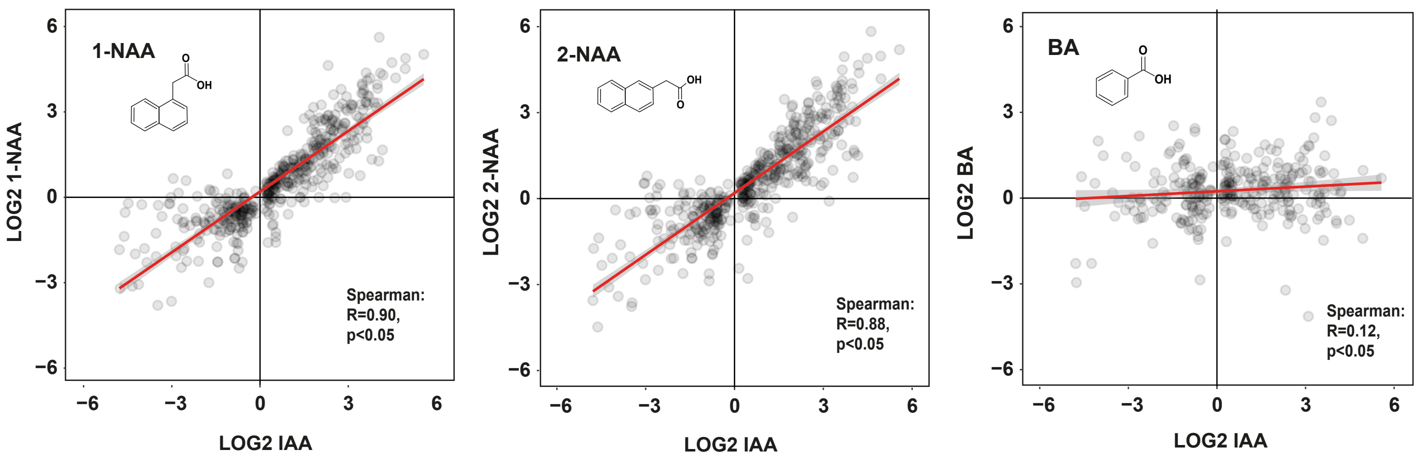

### Figure S2

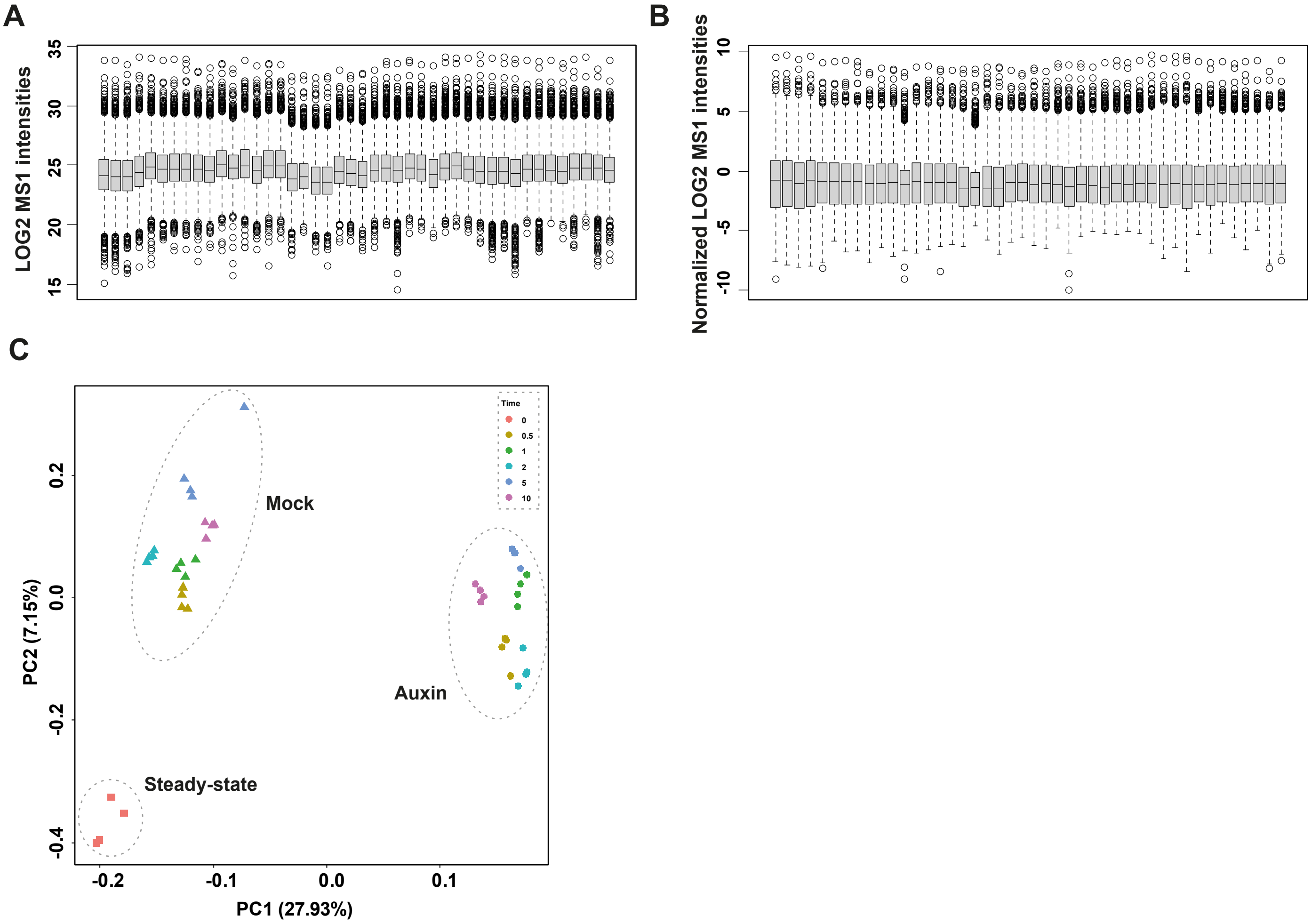

### Figure S3

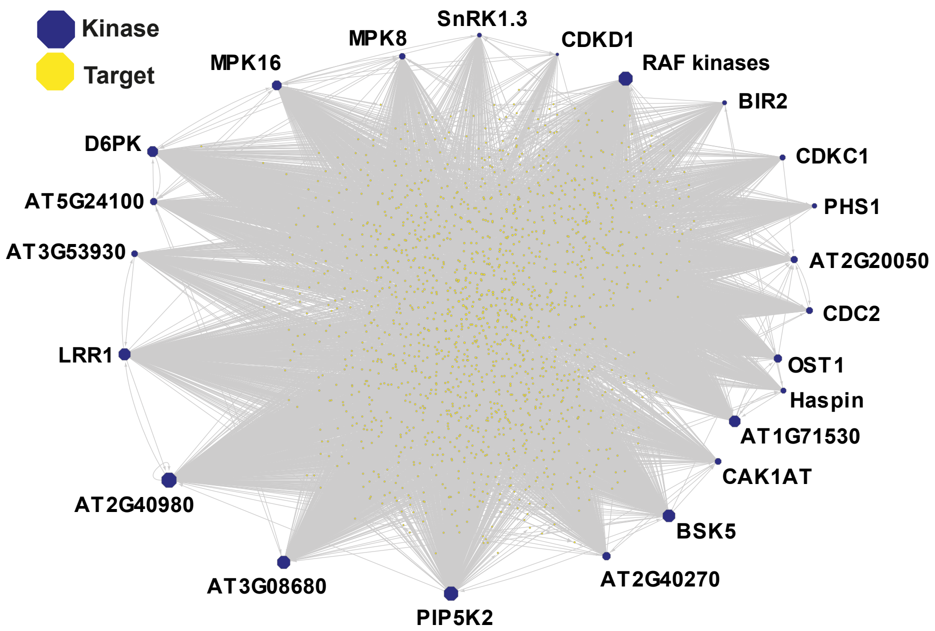

### Figure S4

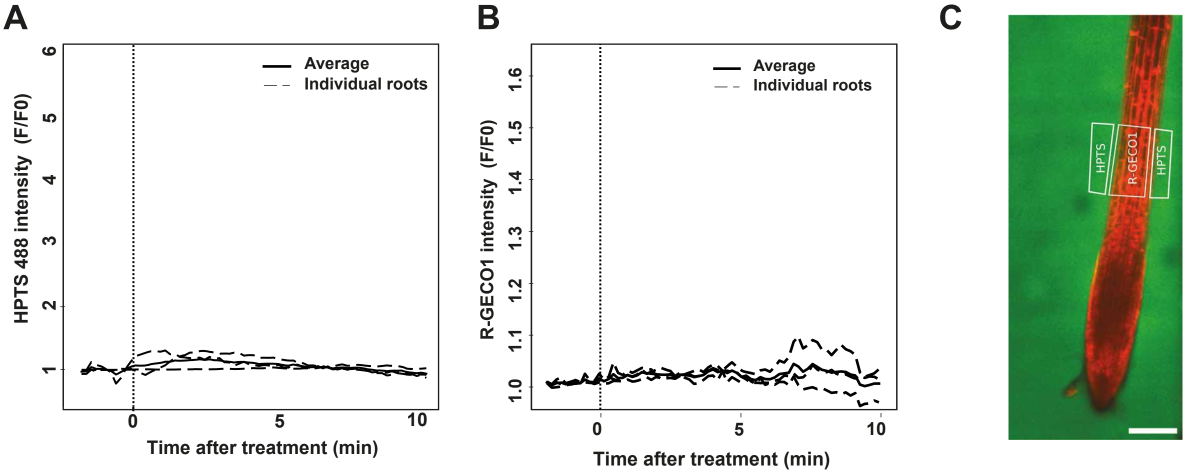
